# A Comprehensive Targeted Panel of 295 Genes: Unveiling Key Disease Initiating and Transformative Biomarkers in Multiple Myeloma

**DOI:** 10.1101/2023.10.28.564536

**Authors:** Vivek Ruhela, Rupin Oberoi, Ritu Gupta, Anubha Gupta

**Author notes:** Corresponding author, (Ritu Gupta). Corresponding author, *Email addresses:* (Anubha Gupta).

## Abstract

Multiple myeloma (MM) is a haematological cancer that evolves from the benign precursor stage termed monoclonal gammopathy of undetermined significance (MGUS). Understanding the pivotal biomarkers, genomic events, and gene interactions distinguishing MM from MGUS can significantly contribute to early detection and an improved understanding of MM’s pathogenesis. This study presents a curated, comprehensive, targeted sequencing panel focusing on 295 MM-relevant genes and employing clinically oriented NGS-targeted sequencing approaches. To identify these genes, an innovative AI-powered attention model, the *Bio-Inspired Graph Network Learning-based Gene-Gene Interaction* (BIO-DGI) model, was devised for identifying *Disease-Initiating* and *Disease-Transformative* genes using the genomic profiles of MM and MGUS samples. The BIO-DGI model leverages gene interactions from nine protein-protein interaction (PPI) networks and analyzes the genomic features from 1154 MM and 61 MGUS samples. The proposed model outperformed baseline machine learning (ML) and deep learning (DL) models on quantitative performance metrics. Additionally, the BIO-DGI model identified the highest number of MM-relevant genes in the post-hoc analysis, demonstrating its superior qualitative performance. Pathway analysis highlighted the significance of top-ranked genes, emphasizing their role in MM-related pathways. Encompassing 9417 coding regions with a length of 2.630 Mb, the 295-gene panel exhibited superior performance, surpassing previously published panels in detecting genomic disease-initiating and disease-transformative events. The panel also revealed highly influential genes and their interactions within MM gene communities. Clinical relevance was confirmed through a two-fold univariate survival analysis, affirming the significance of the proposed gene panel in understanding disease progression. The study’s findings offer crucial insights into essential gene biomarkers and interactions, shaping our understanding of MM pathophysiology.

## 1. Introduction

Multiple Myeloma (MM) is a haematological malignancy characterized by the clonal proliferation of plasma cells within the bone marrow. Monoclonal Gammopathy of Undetermined Significance (MGUS) and MM are both plasma cell disorders, representing distinct stages along the disease progression continuum. MM entails malignant plasma cell proliferation, organ damage, and clinical symptoms, whereas MGUS is a precursor condition with no apparent clinical manifestations. Progression from MGUS to MM occurs at a rate of 1% per year; thus, all MGUS patients do not transition to overt MM during their lifetime [1]. In this context, identifying MGUS individuals likely to progress to MM is crucial for timely intervention and improved outcomes. The clinical distinction between the two conditions primarily relies on tumour load, reflected in monoclonal proteinemia, percentage bone marrow plasma cell infiltration, and end-organ damage [2]. This emphasizes the need to delve into genomic markers and gene-gene interactions to enhance diagnostic accuracy.

The advanced genomic profiling techniques like whole-exome sequencing (WES) and whole-genome sequencing (WGS) enable a thorough examination of genomic aberrations in MM and MGUS. They have proven crucial in identifying key events, including copy number variations (CNVs) and structural variations (SVs) [3]. Numerous MM-related genomic studies have reported significant CNVs, such as del(1p), gain(1q), del(13q), and del(17p), alongside key SVs like translocation involving IgH (e.g. t(4;14), t(11;14), t(14;16), t(8;14)), MYC rearrangement like MYC-IGL, MYC-IGK rearrangements, shedding light on their association with MM prognosis [4, 5, 6, 7, 8, 9, 10]. Moreover, recent studies have highlighted the impact of minor genomic alterations on MM patients’ clinical outcomes [11, 12, 13, 14]. Recently, biallelic alterations in TP53 and DIS3 gene have been reported as high-risk markers in MM [15].

Integrating protein-protein interactions (PPI) with WES and WGS variant profiles can provide crucial insights into genomic biomarkers essential for MGUS to MM progression. Distinguishing MM from MGUS via gene-gene interactions can revolutionize clinical approaches, aiding early detection. This can enable proactive monitoring and intervention for high-risk MGUS patients while sparing low-risk MM cases from aggressive treatments [16].

Several targeted sequencing panels have been devised to comprehensively profile the genomics complexity of MM [17, 18, 19]. These panels encompass critical genomic aberrations related to MM. For instance, a 26-gene panel focused on prevalent mutations in previously published MM-relevant genes [17], but lacked validation for SVs. Similarly, another panel of 182 genes validated for single nucleotide variants (SNVs), CNVs, and specific translocations (related to IGH only) in previously published MM-relevant genes [20]. A more extensive 228-gene panel covered various alterations, including SNVs, CNVs, and translocations involving IgH and MYC genes [18]. In a similar quest for comprehensive genomic profiling of MM, a 47-gene panel was crafted, encompassing dysregulated and frequently mutated genes in MM and those targeted by common therapies, validated for SNVs only [19]. Lastly, the largest gene panel of 465 genes was designed and validated for MM-related SNVs, CNVs, and translocations related to the IGH gene only [21]. However, these panels were designed using only MM samples and hence, lacked markers and interactions distinguishing MM from MGUS that can give insights into MM pathogenesis.

Machine learning (ML) and deep learning (DL) advancements have revolutionized bioinformatics, enabling precise biomarker discovery for early disease detection. Researchers utilize these tools to predict Protein-Protein Interactions (PPIs) and unravel crucial gene interactions in cancer. Notably, models like DeepPPI (Deep neural networks for Protein–Protein Interactions prediction) predict gene interactions based on shared protein descriptors [22]. ML and DL-driven approaches also facilitate inferring semantic similarity of gene ontology terms using PPIs [23, 24, 25]. Despite these strides, no computational model was tailored to identify pivotal biomarkers and gene interactions distinguishing MM from MGUS.

The fusion of genomic mutation profiles with the Protein-Protein Interaction (PPI) data remains an underexplored domain. Graph Deep Learning (GDL) emerges as a potent tool in genomics, promising profound insights from intricate biological graph data structures. Genomic data inherently manifests a graph-like structure, where nodes embody biological entities (genes, proteins) and edges depict relationships. Graph Convolutional Networks (GCNs) play a pivotal role in this realm. GCNs are instrumental in processing and analyzing graph-structured genomic data, where biological entities and their relationships are effectively modelled as a graph, particularly for disease classification [26]. Our previous work seamlessly integrated exonic mutation profiles with PPI to unveil key distinguishing biomarkers of MM and MGUS through the bio-inspired BDL-SP model [27]. We pursued heightened precision in identifying biomarkers and gene interactions by integrating gene interactions from nine diverse PPI databases.

Motivated to bridge this gap, we envisioned a targeted sequencing panel for a thorough genomic profiling of MM, aiming to capture the unique characteristics of MM and MGUS. To address this challenge, we introduced a novel AI-powered attention model: *Bio-inspired Graph network learning based on directed gene-gene interactions (BIO-DGI)*, employing graph network learning to discern differentiating biomarkers and gene-gene interactions in MM and MGUS. In this proposed model, we have integrated bio-inspired learning, utilizing the topological information gathered from nine PPI networks and exomic mutational profiles. This empowered the BIO-DGI model to rank genes and genomic features based on their role in disease progression more efficiently, with fewer graph convolution network layers and multi-head attention modules compared to traditional machine learning (ML) or deep learning (DL) models that relied solely on exomic mutational profiles. The BIO-DGI model also helped us in identifying *Disease-Initiating* and *Disease-Transformative* genes that can aid into understanding MM pathogenesis. This model outperformed exhaustive bench-marking against several baseline ML and DL models, including both quantitative and qualitative evaluations.

We further delved deeper and identified five distinct gene communities using the Leiden algorithm [28] by utilizing the adjacency matrices derived from the five trained BIO-DGI classifiers. This analysis shed light on the influential genes within these communities, quantified through Katz centrality scores [29]. Importantly, we have highlighted genes that were observed to be located in the central position in these gene communities and hence, might be playing a significant role in MM pathogenesis. We analyzed various variant profiles, including SNVs, CNVs, SVs, and Loss of Function (LOF) mutations. This detailed investigation, along with the post-hoc analysis via ShAP (SHapley Additive exPlanations) algorithm [30] that identified top-ranked genes and genomic features, led to the design of a clinically tailored 295-gene panel, aiming for comprehensive genomic profiling of Multiple Myeloma (MM).

Pathway enrichment analysis of these 295 genes revealed enriched MM-related pathways, strongly underscoring the pivotal role of these genes in MM pathogenesis. This discovery, along with the survival analysis corresponding to these genes, underscores the clinical relevance and potential of the targeted sequencing panel designed for comprehensive genomic profiling in MM.

## 2. Materials & Methods

### 2.1. Whole-exome sequencing datasets of MM and MGUS patients

In this study, we included tumour-normal pairs of bone marrow (BM) samples from an MM cohort of 1154 samples and an MGUS cohort of 61 samples sourced from three global repositories of whole-exome sequencing (WES) data. For the MM cohort, 1072 samples were acquired through authorized access to the MMRF dbGaP study (phs000748; phs000348), predominantly comprising American population samples [31]. We also downloaded processed MMRF datasets (version IA12) containing CNVs, SVs, and clinical data from the MMRF Research Gateway. Additionally, we included 82 MM samples from an AIIMS dataset representing the Indian population. In the MGUS cohort, we incorporated 28 MGUS samples from the AIIMS dataset and 33 samples from the European Genome-phenome Archive (EGA) data.

### 2.2. Computational tools and software used for data analysis

We utilized Python computational tools (version 3.9.13) for WES data analysis and visualization. For training all deep learning (DL) models in this study, we employed PyTorch (version 1.12.0+cu113) [32]. Additionally, survival analysis was conducted using the statistical programming language R (version 4.3.1) with the “survival” package [33] (version 3.5.5).

### 2.3. Whole exome sequencing Data: Identification of Significantly altered genes

The WES data obtained from AIIMS and EGA contained the raw fastq files, and the MMRF dataset contained the processed VCF (Variant Call Format) files. The computational workflow for the SNV identification, genomic annotation of SNVs, SNV filtration and grouping, and the identification of significantly altered genes were taken from our previous study [27]. Briefly, raw fastq files from AIIMS and EGA datasets were processed using the standard exome sequencing pipeline [34]. Similar to the MMRF data, the SNVs in AIIMS and EGA WES data were extracted using MuSE [35], Mutect2 [36], VarScan2 [37], and Somatic-Sniper [38] variant callers. The SNVs in AIIMS, EGA and MMRF datasets were annotated using the ANNOVAR database [39]. The annotated SNVs were categorized into three categories based on their functional significance, i.e. synonymous SNVs, non-synonymous SNVs and other SNVs. The benign SNVs were filtered out using FATHMM-XF [40]. Lastly, the annotated SNVs were pooled for MM and MGUS cohorts separately and analyzed for identifying significantly altered genes using the ‘dndscv’ tool [41]. The union of significantly altered genes from all four variant callers for the MM cohort of 1154 samples and the MGUS cohort of 61 samples led to 617 and 362 genes, respectively. Union of these significantly altered genes of MM and MGUS cohorts yielded a set of 824 genes.

For each of these 824 genes, the corresponding protein-protein interactions (PPIs) were extracted from the nine PPI databases (BioGrid [42], Bio-Plex [43], FunCoup [44], HIPPIE [45], HumanNet [46], IntegratedAssociationCorrNet [47], ProteomHD [48], Reactome [49], and STRING [50]) and consolidated to form a merged adjacency matrix. These interactions were then combined to generate a consolidated adjacency matrix, where the criterion for consolidation was the presence of interactions in at least one PPI dataset, denoted as 1 for present and 0 for absent. A total of 26 genes lacked interactions with other significantly altered genes and hence, were excluded resulting in a final set of 798 genes (unioin of 351 genes from the MGUS cohort and 598 genes from the MM cohort) that were used to construct the merged adjacency matrix (Table-S1, Supplementary File-1). Besides the PPI interactions, we extracted 26 genomic features (Table-S3, Supplementary File-3) that included the total number of synonymous SNVs (including UTR3 and UTR5 SNVs), non-synonymous SNVs (including start loss, stop loss, stop gain, exonic, ncRNA-exonic, splicing, frameshift insertion, and frameshift deletion SNVs), and other SNVs (non-frameshift insertion/deletion/substitution, intronic, intergenic, ncRNA-intronic, upstream, downstream, unknown, and ncRNA-splicing SNVs), distributive statistics (median and standard deviation) of variant allele frequency (VAF), allele depth (AD), and four variant conservation scores (GERP [51], PhyloP [52], PhastCons [53], and Mutation assessor [54]). The details of the pre-processing workflow are available in Supplementary File-2.

### 2.4. Proposed Directed Gene-Gene Interaction Learning in Biological Network (BIO-DGI)

This study is aimed towards identifying potential driver genes and uncover essential gene-gene interactions responsible for the progression from MGUS to MM. We introduce an innovative Graph Convolutional Network (GCN)–based attention model named “Bio-inspired network learning based on directed gene-gene interactions” (BIO-DGI). The BIO-DGI model, depicted in Figure-1, harnesses the power of GCN to grasp pivotal gene-gene interactions and forecast potential driver genes.

**Figure 1:**
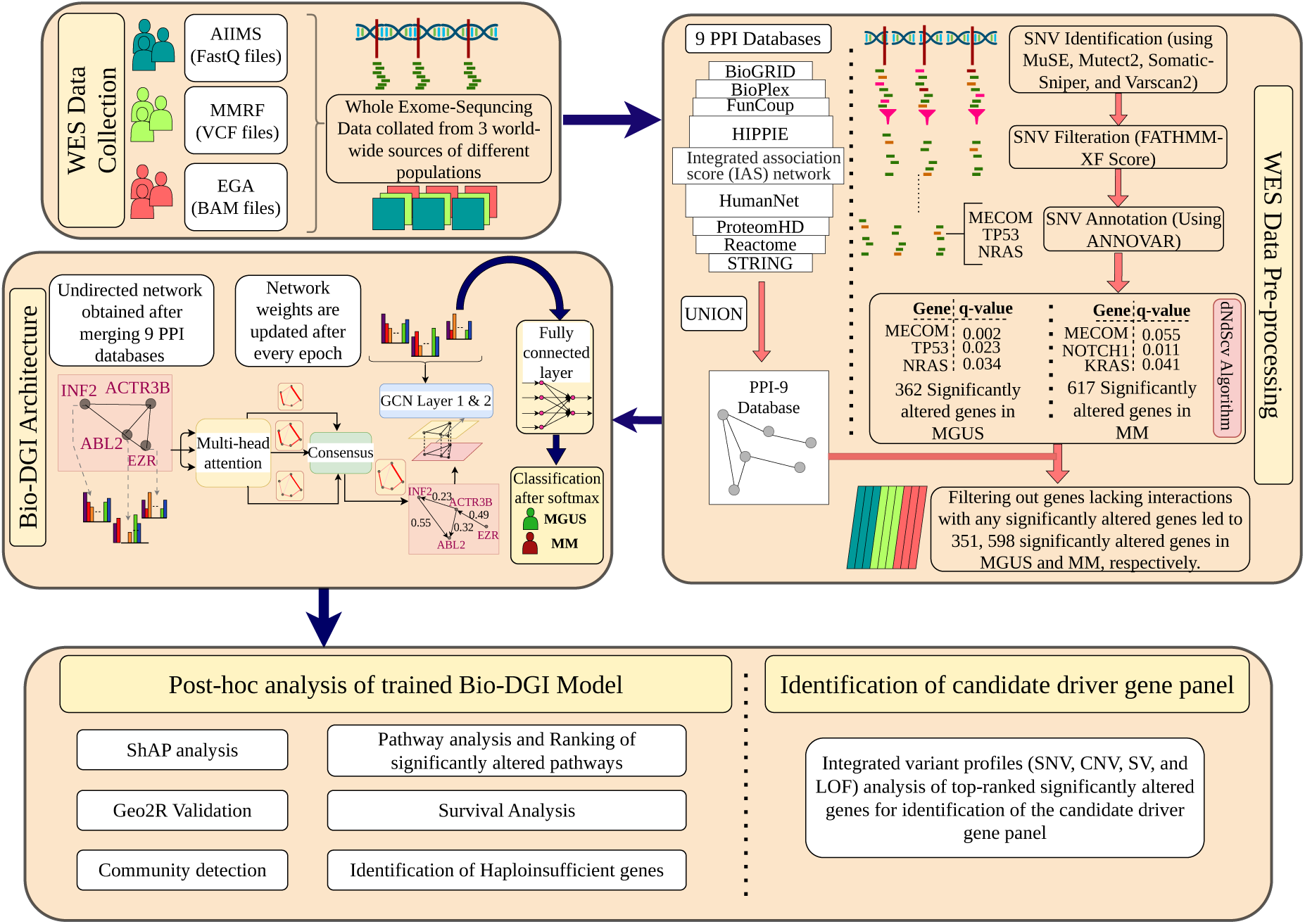
Infographic representation of proposed AI-based bio-inspired BIO-DGI model and post-hoc analysis for identifying pivotal genomic biomarkers that can distinguish MM from MGUS. In the proposed AI-based workflow, the BAM files sourced from EGA and AIIMS datasets, along with VCF files from the MMRF dataset, undergo processing to identify 798 notably altered genes utilizing the dndscv tool (as illustrated in the WES Data pre-processing block). Subsequently, interactions among these 798 genes are elucidated employing PPI networks from nine PPI databases (BioGRID, BioPlex, FunCoup, HIPPIE, IAS network, HumanNet, ProteomHD, Reactome, and STRING). A network is constructed with nodes representing the significantly altered genes and edges denoting interactions obtained after merging interactions from the nine above-mentioned PPI databases. Each node has 26 genomic features (Table-S3, Supplementary File-3) specific to its corresponding gene. The architecture of the BIO-DGI model contains a multihead attention unit and a GCN layer followed by a fully connected layer. The feature matrix and adjacency matrix are provided as input to the BIO-DGI model. The multi-head attention unit in the BIO-DGI model updates the weights of gene interactions in the adjacency matrix, which are then integrated with the sample feature matrix to gain insights on distinguishing biomarkers that can differentiate MM from MGUS. The output of the fully connected layer is converted into the classification probabilities using the softmax activation function. Consequently, the WES data of each subject is analyzed, and feature vectors for all 798 genes are derived. These feature vectors, in conjunction with the subject’s MM/MGUS target class label, constitute the input for supervised training of the GCN. Following learning the BIO-DGI model for distinguishing MGUS from MM, the top genomic features and significantly altered signalling pathways are extracted utilizing the ShAP algorithm and cross-referencing with the Enrichr pathway database.

We supplied two essential inputs to empower the BIO-DGI model: 1) an undirected PPI network adjacency matrix sourced from PPI interaction databases and 2) the feature matrix derived from the data. Two versions of the PPI network adjacency matrix were considered in this study. The first version involved extracting PPI interactions solely from the STRING database, serving as the basis for training the vanilla BIO-DGI model, denoted as BIO-DGI (PPI-STRING). In the second version, we merged the PPI network adjacency matrix from nine distinct PPI databases. This merged adjacency matrix was then utilized for training purposes of the model, referred to as BIO-DGI (PPI9). In both versions of the adjacency matrix, each node corresponded to a significantly altered gene, while the links represented interactions between these genes. Additionally, each node was equipped with a feature vector of length 26, as illustrated in Figure-1. Consequently, the PPI network comprising of 798 significantly altered genes, each associated with a feature vector of length 26, was integrated into the input layer of the BIO-DGI model.

The BIO-DGI model architecture contains 1) a multi-head attention module and 2) a GCN Module. The multi-head attention modules contain three attention units to learn gene-gene interactions, followed by an attention consensus module for taking the consensus of all three attention unit weights to get the updated learned adjacency matrix. The multi-head attention module aimed to learn and update the adjacency matrix to get a weighted PPI adjacency matrix. Similarly, in the GCN module, the input layer is followed by one hidden layer of GCN that is further followed by one fully connected layer of 798 neurons to 2 neurons, giving output through log-softmax activation function for sample class classification (MM vs MGUS).

Our study had 95% MM samples and 5% MGUS samples, which made the data highly imbalanced. Hence, the BIO-DGI model was trained using a cost-sensitive negative log-loss (NLL) function to account for the data imbalance. The BIO-DGI model was trained using a five-fold cross-validation technique that led to the training of five best-performing classifiers. All five classifiers with learned adjacency matrices were saved for further post-hoc analysis. We used the ShAP algorithm for post-hoc analysis of BIO-DGI model classifiers to get top-performing genes and genomic features that were further used for pathway enrichment analysis, gene-community identification and candidate driver gene panel. The setting of layers, hyperparameters used to train the BIO-DGI model, and mathematical description of the BIO-DGI model are available in Supplementary File-2.

### 2.5. Quantitative benchmarking of BIO-DGI model with traditional Machine learning classifiers

In our quantitative benchmarking analysis, we conducted a comprehensive comparison of the BIO-DGI (PPI9) model involving three key performance metrics: balanced accuracy, area under the curve (AUC), and area under the precision and recall curve (AUPRC). This evaluation encompassed the five-fold cross-validation of six established baseline cost-sensitive machine learning models: random forest, decision tree, logistic regression, XGBoost, CatBoost, and SVM from the scikit-learn library [55]. Further, we also included two cost-sensitive DL models: BDL-SP and BIO-DGI (PPISTRING) models for quantitative benchmarking. We incorporated a tailored cost-sensitive loss function to enhance the models’ sensitivity to class imbalance. This function implements weighted penalization for sample misclassifications, with the weighting being directly proportional to the class imbalance ratio. This strategic implementation of weighted penalization ensures unbiased learning outcomes for major and minor classes, fostering a more equitable predictive capability.

### 2.6. Identification of gene communities using learned adjacency matrices of BIO-DGI

We employed a five-fold cross-validation training strategy to our proposed BIO-DGI (PPI9) model. Subsequent to training the model, we retained the learned adjacency matrix from each classifier, yielding five distinct learned adjacency matrices. We individually identified gene communities from these adjacency matrices using the Leiden algorithm. This process yielded 5, 5, 6, 5, and 6 gene communities across the learned adjacency matrices. From the communities extracted from each adjacency matrix, the top 3 communities were selected based on the number of OGs, TSGs, ODGs, and AGs. Consequently, we retained the weights of only those genes in the adjacency matrix that were a part of these top 3 communities, while the links of rest of the genes were dropped by assigning zero weight. We called this modified adjacency matrix as the *Gene Community Adjacency* (GCA) matrix. This process was carried out for all the five learned adjacency matrices (one matrix learned from the training of each fold classifier). Next, we computed the mean GCA matrix by calculating the mean weight of a gene across all GCAs. Lastly, we applied the Leiden algorithm for community detection on the mean GCA matrix to obtain the consolidated gene communities.

### 2.7. Qualitative application-aware post-hoc benchmarking of BIO-DGI model

The ShAP (SHapley Additive exPlanations) algorithm is a powerful tool for gauging the significance of attributes in a model’s predictions. It achieves this by assigning scores to attributes based on their individual contributions. In this context, ShAP played a pivotal role in enhancing the post-hoc explainability of the BIO-DGI (PPI9) model. This process unearthed the most influential genomic features and the genes that experienced significant alterations, both at the cohort level (MGUS or MM) and at the level of individual samples. The ShAP algorithm was applied to each trained classifier obtained after a rigorous five-fold cross validation was carried out during the model’s training phase. This enabled the identification of significant genomic attributes (genes and features) for every sample. It is important to note that a ShAP score can have both positive and negative values, wherein a positive ShAP score for a specific attribute highlights its contribution to the model’s prediction for the MM class (positive class). Conversely, a negative score indicates its role in the model’s prediction for the MGUS class (negative class). Consequently, the magnitude of the ShAP score directly correlates with the attribute’s impact on the model’s positive class outcome. Furthermore, the extraction of ShAP interpretability was limited to samples correctly predicted by at least one of the five classifiers. This approach ensured a robust basis for deriving insights through ShAP analysis.

We estimated the best ShAP scores on a per-sample basis: 1) for all 798 significantly altered genes and 2) for all 26 genomic features. To this end, for each sample in the MM and MGUS cohort, class predictions were taken from all the five trained classifiers of the BIO-DGI (PPI9) model. Next, the inference of the ShAP algorithm was taken for only the classifiers that made correct predictions for that sample. ShAP scores were collected both at the classifier and sample levels for all genomic attributes. Next, for all the significantly altered genes, the ShAP scores of the 26 genomic features were grouped by their positive and negative signs. The best ShAP score for each gene was determined by comparing the absolute values of these grouped scores, considering the highest absolute value among all classifiers as the best possible score. Similarly, for each genomic feature, the ShAP scores of all 798 genes were grouped and assessed in a similar manner, resulting in the best ShAP score. Following this process, the most highly ranked genes and genomic attributes were identified at both the cohort and sample levels.

We extended our analysis by comparing the BIO-DGI (PPI9) model’s top-ranked significantly altered genes with those reported in previous studies, aiming to identify genes previously reported to be associated with disease progression or suppression. We validated and analyzed our model using information from multiple databases such as OncoKB [56], IntoGen [57], COSMIC [58], and TargetDB [59] at the gene level. For model validation, we extracted 1064 cancer genes from the OncoKB database for oncogenes and tumour-suppressor genes. From the COSMIC database, we utilized 318 oncogenes and 320 tumor-suppressor genes.

We utilized the IntoGen database (https://www.intogen.org/) and MM-related studies [12, 60] to compile a catalogue of MM driver genes. Additionally, 180 actionable genes from COSMIC and 135 from TargetDB helped infer actionable genes. Using the above information, we regrouped our top-ranked significantly altered genes into Oncogenes (OGs), Tumor-Suppressor genes (TSGs), Onco-driver genes (ODGs), and Actionable genes (AGs). This comprehensive approach facilitated a thorough exploration of genomic features in the post-hoc interpretability analysis of the BIO-DGI (PPI9) model, providing valuable insights into their roles in disease contexts. Furthermore, we introduced another way of understanding the disease pathogenesis by assessing whether a gene was observed to be significantly altered in MM or MGUS or both. Genes found to be significantly mutated in only MM (and not in MGUS) were designated as *Disease-Transformative* genes, while those significantly altered in both MM and MGUS were labelled as *Disease-Initiating* genes. This new way of categorization deepened our understanding of these genes, shedding light on their biological functions and specific relevance to MM and MGUS. This comprehensive approach facilitated a thorough exploration of genomic features in the post-hoc interpretability analysis of the BIO-DGI (PPI9) model, providing valuable insights into their roles in the context of disease.

### 2.8. Identification of CNVs, SVs and LOFs for top 500 significantly altered genes

Our exploration into significantly altered genes underwent expansion to encompass a broader array of genomic profiles, including copy number variants, structural variants, and loss-of-function events. This extended analysis allowed us to delve into the impact of these variants at the gene level, shedding light on their influence on disease progression. For the MMRF dataset, we leveraged the segment data obtained from MMRF CoMMpass to identify copy number variants (CNVs) using the CNVkit [61] tool. To ensure consistency in our CNV identification workflow, we applied CNVkit to detect CNVs in the WES samples of both AIIMS and EGA datasets. Within this framework, we filtered out genes with a copy number (CN) value of 2 across all samples, focusing on genes with CN values that deviated from 2. Next, we utilized all the structural variants identified in WGS samples from the MMRF dataset (i) by the Translational Genomics Research Institute (TGen) through their in-house SV identification workflow and (ii) by the Delly tool [62]. Our analysis centred on significantly altered genes ranked in the top 500, whose genomic regions were affected by structural variants, spanning insertions, inversions, deletions, duplications, and translocations.

Furthermore, our investigation extended to encompass genes marked by loss-of-function aberrations. Loss-of-function refers to a disruption in a gene’s normal functioning, hindering the production of the typical gene product or rendering it ineffective. We assessed every transcript of a gene in a sample to ascertain if it satisfied any of the following conditions: deletion of over half of the coding sequence, deletion of the start codon, deletion of the first exon, deletion of a splice signal, or deletion causing a frameshift, it was considered to exhibit loss-of-function [63]. If at least one condition was satisfied by all the transcripts of a gene in a sample, that sample was labelled to exhibit LoF in that gene [63]. This evaluation was conducted for all the MM and MGUS samples to identify genes featuring loss-of-function.

### 2.9. Geo2R Validation of top 500 significantly altered genes

We conducted a thorough validation using the Geo2R tool [64] to validate 295 genes against previously published studies focused on multiple myeloma (MM). We utilized a total of eleven micro-array and miRNA data-based MM studies for the validation [65, 66, 67, 68, 69, 70, 71, 72, 73, 74, 75, 76, 77]. To ensure rigorous assessment, we exclusively considered genes that displayed significant dysregulation and maintained an adjusted *p*-value *≤* 0.05 in each validation instance.

### 2.10. Workflow for the design of targeted sequencing gene panel

We devised an innovative workflow to identify potential driver genes for designing the targeted sequencing gene panel for MM. This extensive process integrated various genomic profiles, including single nucleotide variations (SNVs), copy number variations (CNVs), structural variations (SVs), and loss-of-function (LoF) mutations, alongside validated datasets from Geo2R. Initially, we focused on the top 500 genes, extracting relevant data from their SNV, CNV, SV, and LoF profiles. Following this, we implemented a rigorous filtering criteria to identify pivotal genes within each variant profile. This involved generating box plots for the profiling feature across all samples and subsequently applying filtering criteria based on statistical analysis. For instance, for LOF, we made the box plot of number of samples having LoF for every gene of the MM cohort and retained only those genes that were found to have LOF in the number of samples greater than the 3rd quartile of this box plot. Similar criteria were applied on the other variant profiles, detailed in Figure-2. The resulting gene list was consolidated, retaining those meeting criteria in at least one variant profile and had at least one dataset validation in Geo2R analysis. with Geo2R validation. We specifically retained disease-transformative and disease-initiating genes, dropping those genes that were significant only in monoclonal gammopathy of undetermined significance (MGUS) but not in the MM cohort, yielding a final list of 282 genes.

**Figure 2:**
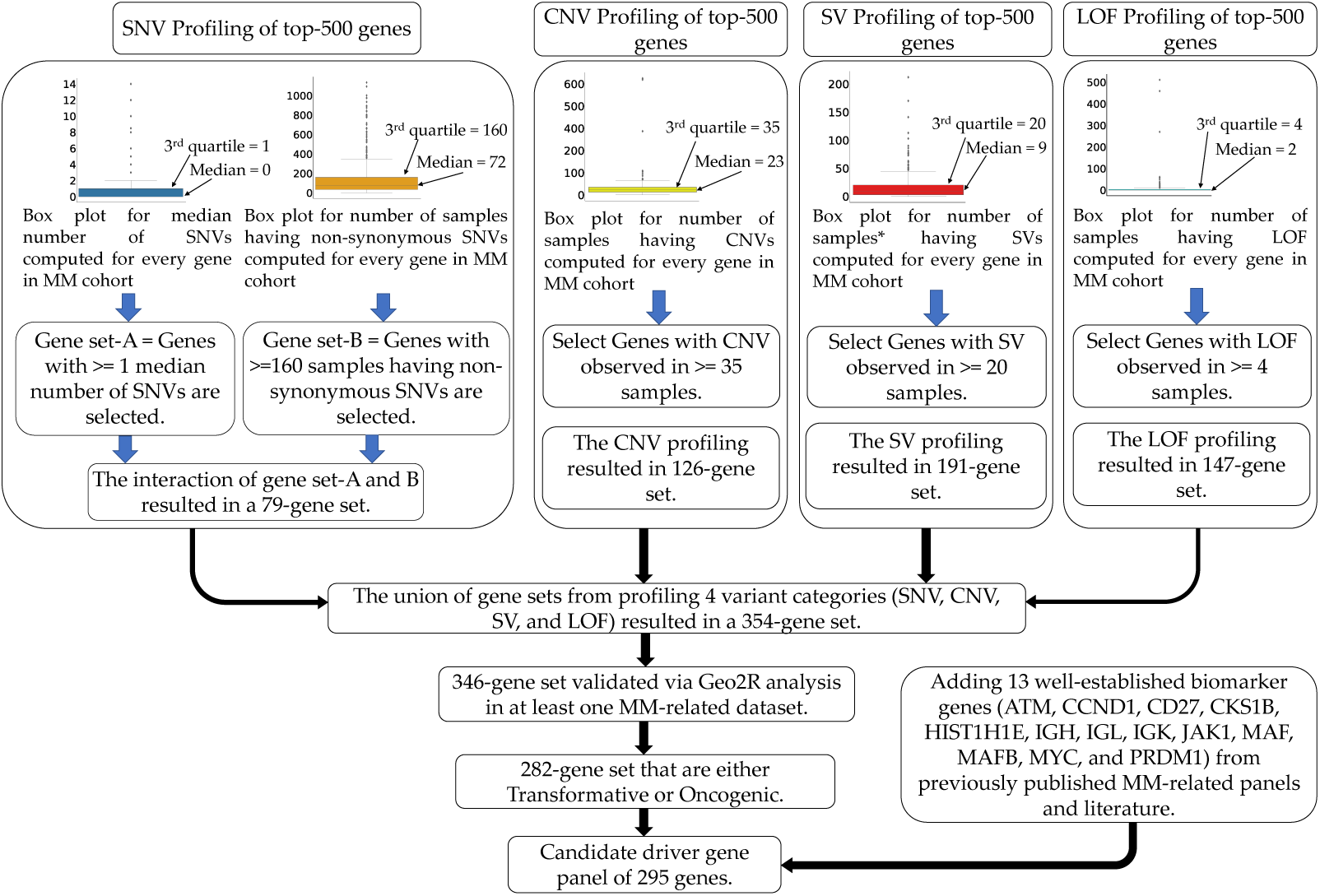
Workow for the Identification of Biomarker Genes for the Proposed 295-Gene Panel. The workow integrates variant profiles, including SNVs, CNVs, SVs, and LOF, to discern genes relevant to MM. The combination of these profiles yielded a gene set of 354 candidates, which was further refined to 346 genes after Geo2R validation. From this set, disease-initiating genes (significantly altered in both MM and MGUS) and disease-transformative genes (significantly altered only in MM) were selected for inclusion in the targeted sequencing panel, resulting in a final list of 282 genes. Additionally, thirteen well-established biomarker genes from previously published MM-related panels and literature were incorporated, culminating in the completion of the final 295-gene panel.

Since the analysis originated from SNVs extracted from the WES data and certain key MM genes like IGH and MYC, known for translocations in MM, were not initially visible, we incorporated twelve additional well-established MM biomarkers (ATM, CCND1, IGH, IGL, IGK, CKS1B, HIST1H1E, JAK1, MAF (or c-MAF), MAFB, MYC, and PRDM1), yielding our panel of 295 genes. These genes, recognized for CNV or SV profiles, were reported in at least two previously published MM-related panels studied in this work. The detailed workflow for potential driver gene identification is presented in Figure-2. Lastly, we assessed the major molecular aberrations for each gene in this panel of 295 genes and examined coding regions and genomic locations for altered regions using the UCSC Genome database [78] to understand the genomic spectrum of MM.

### 2.11. Identification of CNVs, SVs and LOFs of the proposed gene panel and Geo2R Validation

Building upon the genomic profile analysis of CNVs, SVs, and LOFs in the top 500 genes presented in Section 2.8, we further investigated the proposed 295-gene panel by examining its CNV, SV, and LOF landscape. This provided deeper insights into the panel’s suitability for MM diagnosis and risk stratification. Additionally, we leveraged Geo2R to validate the panel’s relevance against existing MM-related studies, bolstering its potential clinical utility.

### 2.12. Workflow for the comprehensive survival analysis of the proposed gene panel

Building upon the targeted sequencing gene panel designed in Section-2.10, we conducted a novel survival analysis to investigate the impact of gene variant profiles on patient survival in Multiple Myeloma (MM). Employing two distinct approaches outlined below, we sought to discern the genes that exert a statistically significant impact on the survival outcomes of these patients.

In the first approach, univariate survival analysis was conducted for all 295 genes individually, considering each variant profile (SNV, CNV, SV, and LOF) as a singular prognostic factor. For the SNV profile, we utilized the total count of (non-synonymous + other) single nucleotide variants (SNVs) as the prognostic factor in the univariate survival analysis. Analogously, for CNV, SV, and LOF profiles, we constructed categorical vectors (yes/no) indicating the presence or absence of copy number variations, structural variations, and loss-of-function mutations in the multiple myeloma (MM) sample for each gene. Subsequently, we performed univariate survival analysis for each variant profile separately. Genes with a *p*-value *≤* 0.05 in the univariate survival analysis for individual variant profiles were retained.

In a parallel vein, the second approach amalgamated all four variant profiles for each gene, with an aim to elucidate the cumulative impact of gene variant profiles on clinical outcomes. Factor Analysis of Mixed Data (FAMD) [79] was employed for dimensionality reduction in this process. Subsequently, univariate survival analysis were executed on each of the 295 genes in the panel, utilizing the first FAMD component as the prognostic factor. Genes with a *p*-value *≤* 0.05 in the univariate survival analysis of the first FAMD component were retained. Finally, we considered the union of genes identified as clinically relevant (*p*-value *≤* 0.05) through the aforementioned two approaches.

### 2.13. Identification of significantly altered pathways and pathway ranking using the gene panel

Out of 295 genes, the noteworthy 282 genes highlighted by the BIO-DGI (PPI9) model as instrumental in distinguishing MM from MGUS and included in the 295-genes panel via the workflow of Figure-2 were cross-referenced with the significant gene lists derived separately for MM and MGUS cohorts using the dndscv tool. The MM cohort genes were specifically employed in the pathway analysis for MM, while the MGUS cohort genes were utilized for MGUS pathway analysis. Notably, thirteen genes included in the 295-gene panel due to their association with translocations specific to MM disease were integrated with the gene list of the MM cohort for the aforementioned pathway analyses.

To further elucidate the functional implications, we employed the ‘Enrichr gene set enrichment analysis web server’ [80, 81, 82], facilitating the identification of KEGG and Reactome pathways associated with our proposed gene panel. Subsequent ranking of significantly altered pathways in the MM and MGUS cohorts, based on their adjusted *p*-values, provided a comprehensive insight into the primary pathways undergoing substantial alterations due to genomic aberrations in the significantly altered genes.

### 2.14. Identification of Haploinsufficient genes of the gene panel

To assess the likelihood of genes exhibiting haploinsufficiency, we draw upon two previously published haploinsufficiency prediction scores: the genome-wide haploinsufficiency score (GHIS) [83] and the DECIPHER score [63]. The DECIPHER score amalgamates patient genomic data, evolutionary profiles, and functional and network properties to predict the likelihood of haploin-sufficiency. Meanwhile, the GHIS score draws from diverse large-scale datasets, encompassing gene co-expression and genetic variation in over 6000 human exomes. These comprehensive methods enhance identifying haplo-insufficient genes, revealing their crucial role in diseases. This deepens our understanding of genes that lack proper function when only one copy is present.

## 3. Results

### 3.1. Cohort Description

In this comprehensive study, we analyzed two distinct cohorts related to MM and MGUS, encompassing a total of 1154 MM samples and 61 MGUS samples sourced from three globally recognized datasets: AIIMS, EGA, and MMRF. Specifically, within the MM cohort, we examined 1072 samples from the MMRF dataset and 82 samples from the AIIMS dataset. Additionally, in the MGUS cohort, we examined 28 samples from the AIIMS repository and 33 from the EGA repository. Augmenting our analysis, we incorporated crucial clinical data, including overall survival (OS) time and event data for MM samples retrieved from the MMRF and AIIMS datasets. This enabled a thorough exploration of the clinical relevance of the proposed targeted sequencing panel, underlining the significance of our findings.

### 3.2. Identification of Significantly altered genes

We employed the DNDSCV tool, as illustrated in the pre-processing block of Figure-1, to discern genes exhibiting significant alterations within the MM and MGUS cohorts. A total of 598 and 351 significantly altered genes were identified in the MM and MGUS cohorts, respectively. Notably, 151 genes were found to be shared between the MM and MGUS cohorts. Subsequently, we pursued the inference of pivotal genes and gene-gene interactions vital for discriminating MM from MGUS, leveraging our innovative graph-based BIO-DGI (PPI9) model.

### 3.3. Benchmarking of proposed BIO-DGI (PPI9) model

Employing our AI-driven BIO-DGI workflow (depicted in Figure-1), we trained the BIO-DGI (PPI9) model using 5-fold cross-validation and compared its performance with six standard cost-sensitive machine learning and two deep learning models. Remarkably, the proposed BIO-DGI (PPI9) model showcased superior performance in terms of balanced accuracy and AUPRC (area under the precision-recall curve). Specifically, the BIO-DGI (PPI9) model achieved the highest balanced accuracy of 96.7%. Following closely, the BDL-SP model attained a balanced accuracy of 96.26%, and the cost-sensitive Catboost (CS-Cat) model achieved the third-best performance with a balanced accuracy of 96.09%. The BIO-DGI (PPI9) model also outperformed other models in AUPRC, securing the highest AUPRC score of 0.93, while the AUPRC score for BDL-SP and CS-Cat models stood at 0.92 and 0.9, respectively. Notably, the BIO-DGI (PPI9) model correctly identified 1099 out of 1154 MM samples and 60 out of 61 MGUS samples, showcasing its superior performance.

These results affirm that, quantitatively, the BIO-DGI (PPI9) model performed superior with the BDL-SP model being the second best. For a comprehensive understanding of quantitative performance, refer to Figure-3 (A) and (B) for balanced accuracy and AUPRC scores, confusion matrices, and AUPRC curves, respectively.

**Figure 3:**
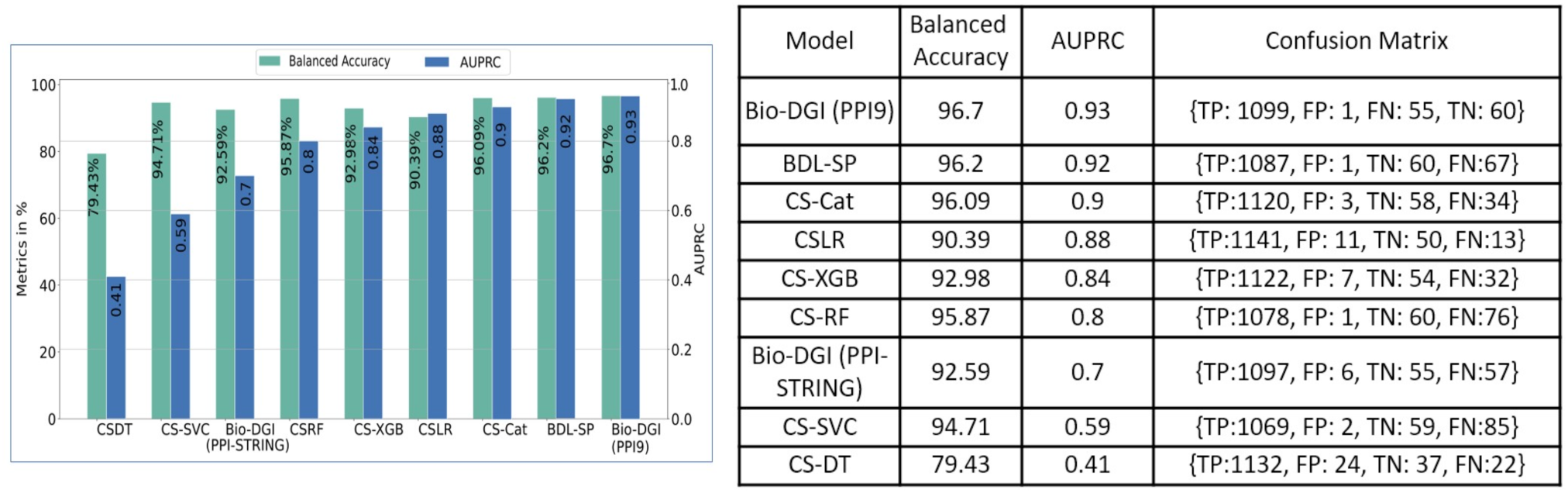
Quantitative benchmarking of proposed BIO-DGI(PPI9) model. (A) Comparison of balanced accuracy and AUPRC score of BIO-DGI(PPI9) model with other baseline ML and DL models, and (B) Confusion matrix of top-performing models including BIO-DGI (PPI-STRING) model. Notations: Cost-Sensitive Decision Tree (CS-DT); Cost-Sensitive Support Vector Classifier (CS-SVC); Cost-Sensitive Random Forest (CS-RF); Cost-Sensitive XGBoost (CS-XGB); Cost-Sensitive Logistic Regression (CS-LR); Cost-Sensitive CatBoost (CS-Cat).

Given the marginal difference in the balanced accuracy and AUPRC performance metrics among the top three models (BIO-DGI (PPI9), BDL-SP, and CS-Cat), we conducted post-hoc interpretability benchmarking by applying ShAP algorithm to identify the top-ranked genes for each of the top three performing models. Subsequently, we analyzed these genes to identify previously reported oncogenes (OG), tumour-suppressor genes (TSG), both oncogenes and driver genes (ODG), and actionable genes (AG). Out of the total 798 genes, we identified 31 OGs (including *ABL2, BIRC6, FUBP1, IRS1*), 43 TSGs (including *APC, ARID1B, CYLD, PABPC1, ZFHX3*), 10 ODGs (including *BRAF, FGFR3, TP53, TRRAP*), and 19 AGs (including *ARID2, BRD4, MITF, NF1, TYRO3*) (Table-1A).

**Table 1:**
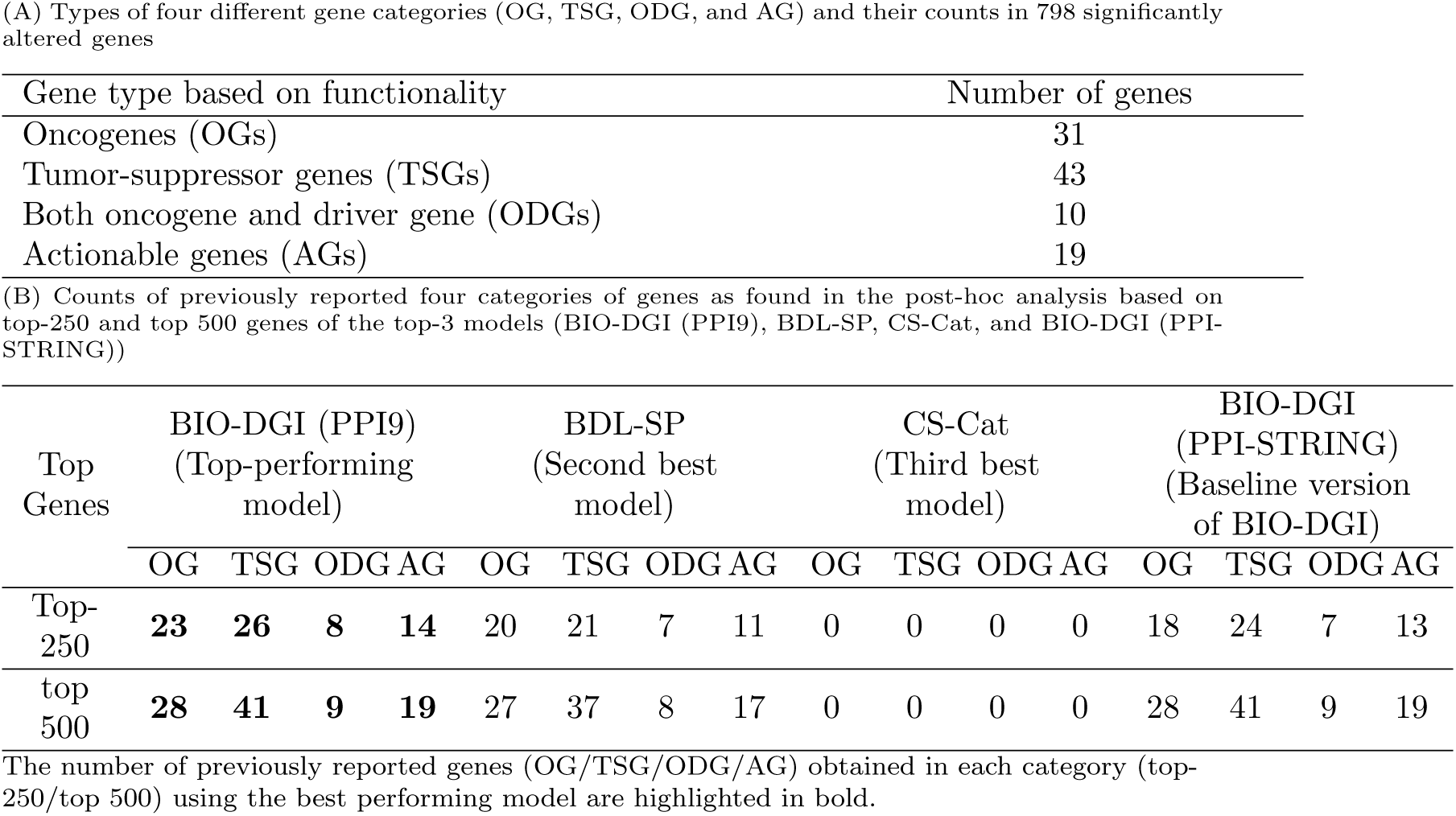
Number of previously reported genes present in 798 significantly altered genes and qualitative benchmarking of top-performing models.

Our analysis revealed that the proposed BIO-DGI (PPI9) model exhibited the highest count of identified oncogenes (OG), tumor-suppressor genes (TSG), both oncogene and driver genes (ODG), and actionable genes (AG) in both the top-250 and top 500 gene lists (Table-1B). Specifically, the BIO-DGI (PPI9) model detected 23 and 28 oncogenes in the top 250 and top 500 gene list, respectively. Of the 43 known TSGs, the BIO-DGI (PPI9) model identified 26 genes in the top 250 and 41 in the top 500 gene lists. Of the 10 known ODGs, the BIO-DGI (PPI9) model identified 8 genes in top 250 and and 9 genes in top 500 gene lists. Lastly, of the 19 known AGs, the BIO-DGI (PPI9) model identified 14 genes in top 250 and and 19 genes in top 500 gene list.

We have considered only those genes in the top-250 or top 500 gene list that have a non-zero ShAP score in the post-hoc explainability analysis. The total counts of previously reported genes as found in the top-250 and top 500 genes of the top-four models (BIO-DGI(PPI9), BDL-SP, CS-Cat, BIO-DGI (PPI-STRING)) is shown in Table-1B. Furthermore, the lists of top 500 genes obtained using Bio-DGI (PPI9) and the previously reported genes ranked within the top 250 and top 500 by the top-performing models are outlined in Table-S2, Supplementary File-1 and Table-2.

**Table 2:**
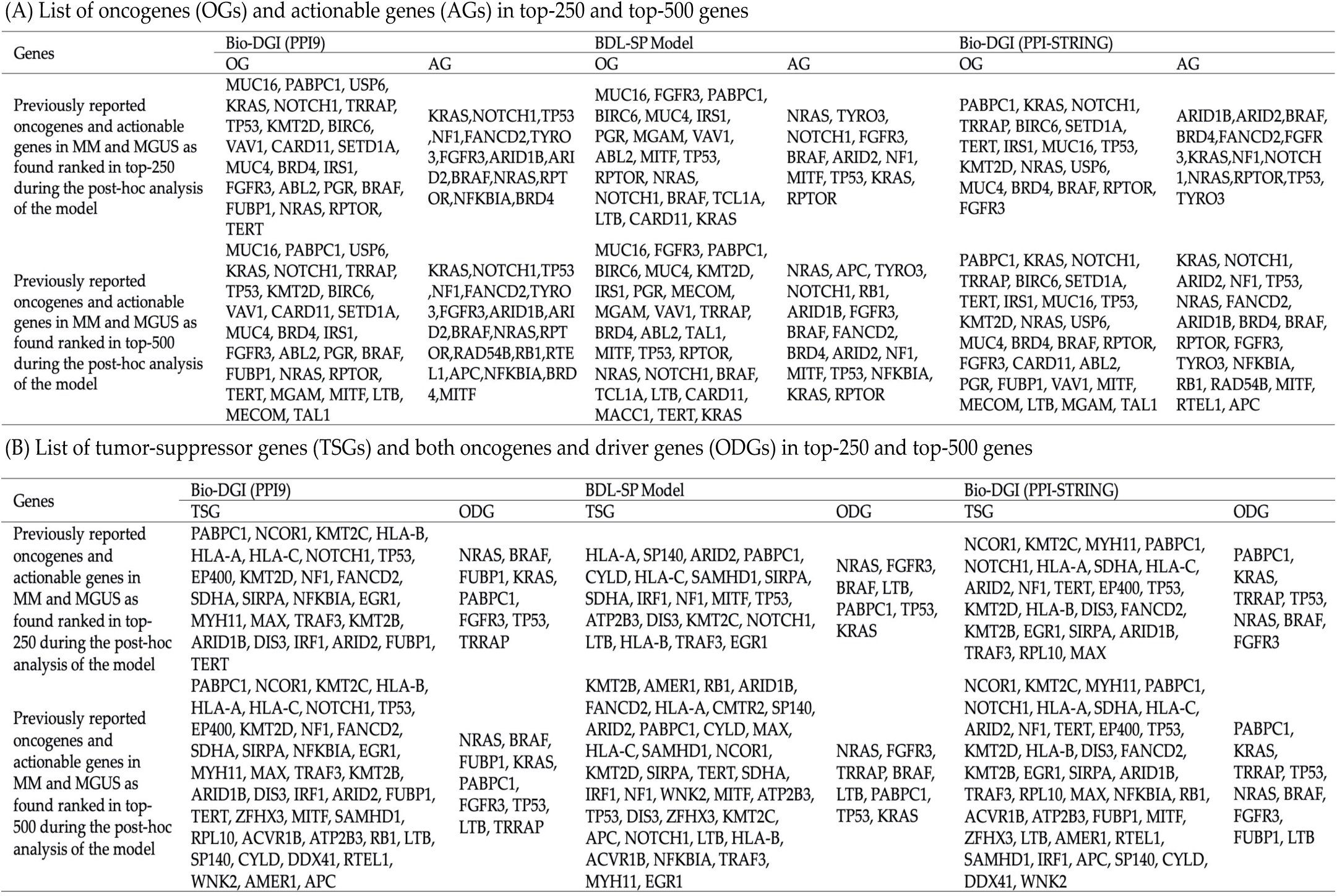
List of 4 categories of previously reported genes as found in the post-hoc analysis based on top-250 and top 500 genes of the top-3 models (BIO-DGI (PPI9), BDL-SP, and BIO-DGI (PPI-STRING))

Given the BIO-DGI (PPI9) model’s superior identification of previously reported OGs, TSGs, ODGs, and AGs, it also stands out as the best-performing model in the post-hoc analysis and was subsequently used to infer the top significantly altered genes, gene-gene interactions, genomic features, and altered signalling pathways critical for distinguishing MM from MGUS. This analysis underscores the importance of model interpretability within the application domain, particularly, when similar quantitative results are obtained with different machine learning models.

### 3.4. Interpretability of BIO-DGI (PPI9) model using ShAP algorithm

We utilized the ShAP algorithm for post-hoc model explainability and rank genomic attributes based on their influence on the model prediction. Each genomic attribute received a ShAP score, representing its contribution to each class (MM/MGUS). Subsequently, the attributes were ranked at the cohort level (MM versus MGUS) accordingly. This ShAP analysis provided post-hoc explainability of the trained model, following a methodology akin to that outlined in [27], enabling the ranking of genes and genomic features at both cohort and sample levels.

By evaluating the ShAP scores assigned to each gene, we identified *MUC6, LILRA1*, and *LILRB1* as the top three genes in MM and MGUS samples among the 798 significantly altered genes. Furthermore, several previously reported oncogenes (e.g., *MUC16, USP6, BIRC6, VAV1*), tumor-suppressor genes (e.g., *EP400, HLA-B/C, SDHA, MYH11*), both oncogenes and driver genes (e.g., *PABPC1, KRAS, TRRAP, TP53, FGFR3, BRAF*), and actionable genes (e.g., *NOTCH1, FANCD2, TYRO3, ARID1B*) were highlighted as top-ranked genes.

Similarly, we ranked genomic features based on their impact on the model’s prediction using their ShAP scores. In our model training for BIO-DGI (PPI9), a set of 26 genomic features was employed. Notably, the PhyloP score of non-synonymous SNVs, allele depth of synonymous SNVs, and the total number of other SNVs (that included non-frameshift insertion/deletion/substitution, intronic, intergenic, ncRNA-intronic, upstream, downstream, unknown, and ncRNA-splicing SNVs) emerged as the top three genomic features. Figure-4 presents the beeswarm plot illustrating the

**Figure 4:**
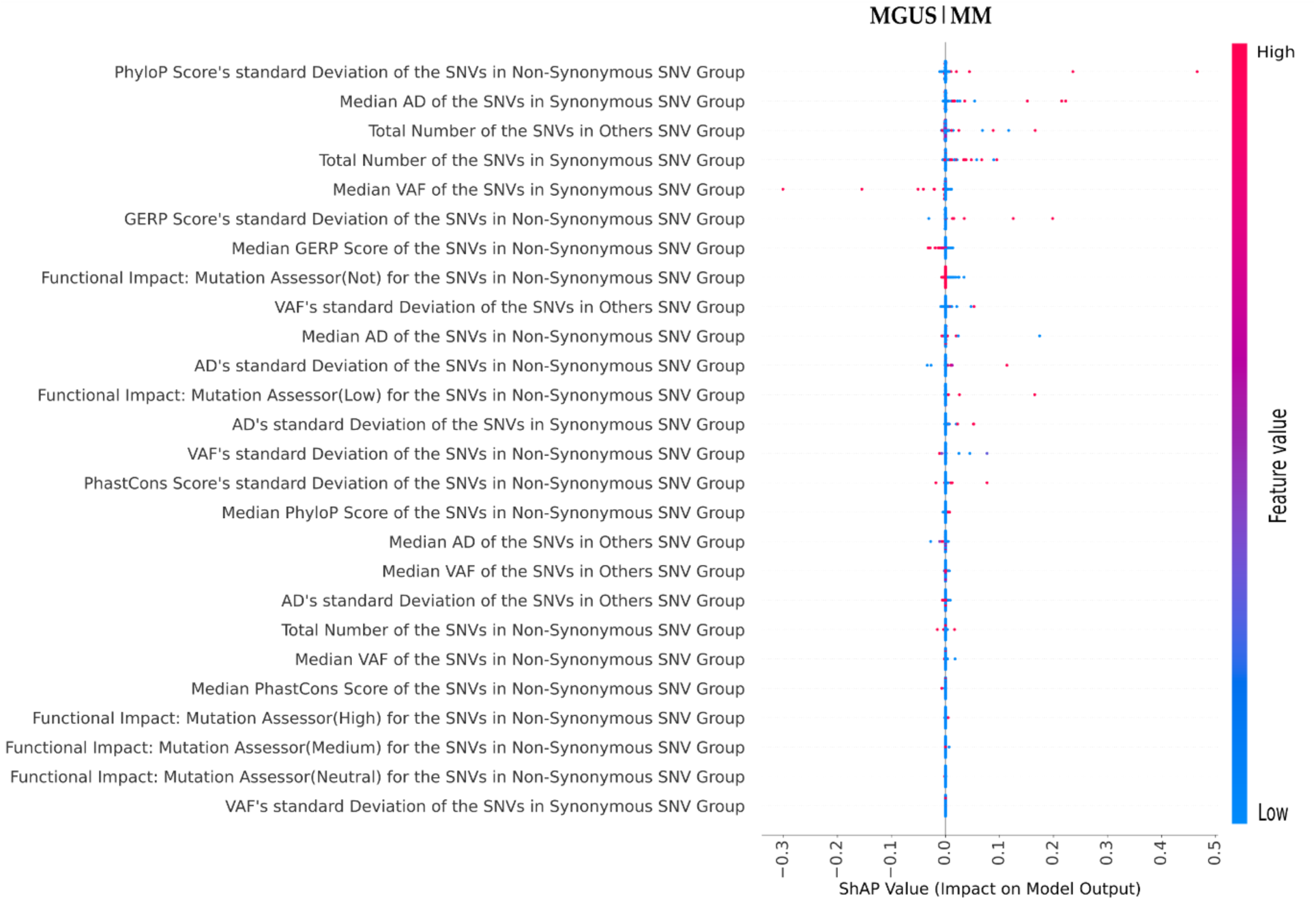
Genomic Feature Ranking using the ShAP Algorithm in MM and MGUS based on post-hoc explainability by the BIO-DGI model. Genomic features are ranked according to their ShAP scores. A positive ShAP score indicates the feature’s contribution to MM, while a negative score represents its contribution to MGUS. Each dot in the scatter plot represents a sample colour-coded to reflect genomic feature values—dark blue for low and red for high values.

### 3.5. Analysis of CNVs, SVs and LOF of top-500 genes in MM

In addition to analyzing SNV profile, we comprehensively investigated CNVs, SVs, and LOF of the top 500 genes of the MM cohort. CNV identification was performed using CNVkit on AIIMS MM samples and on exome segment data from MMRF CoMMpass for MMRF samples. Processed SV data from MMRF CoMMpass was utilized to identify key SVs in MM and 295-genes panel designing. For identifying genes with LOF within a sample, we employed established criteria to evaluate disruptions in gene transcripts due to deletion of essential coding segments, exons, splice signals, or frameshift-inducing deletions [63]. We studied both CNVs and SNVs to identify genes with LOF within each sample.

CNVs, SVs, and LOF analysis of the top 500 genes revealed crucial molecular aberrations in MM. Chromosome-wise distribution analysis indicated that chr19 (19%), chr1 (17%), chr6 (8.6%), and chr14 (7.1%) were notably affected by CNVs (Figure-5(A)). Similarly, chr1 (12.6%), chr6 (9.9%), chr12 (5.3%), and chr14 (5%) showed prominent SV involvement (Figure-5(B)), while chr19 (20%), chr1 (19.9%), chrX (13%), and chr14 (11.8%) were most affected by LOF (Figure-5(C)). The majority of CNVs were gains (58.3%) and deletions (17%) (Figure-5(D)), while inversions (65%) and translocations (13.1%) dominated the SV landscape (Figure-5(E)). Notable chromosomes impacted by inversion SV included chr1, chr3, chr2, and chr7 (Figure-5(F)), and translocations mainly affected chr7, chr21, chr1, and chr14 (Figure-5(G)). The distribution of CNV and SV types within each chromosome highlighted their relative abundance (Figure-5(H) and Figure-5(I)).

**Figure 5:**
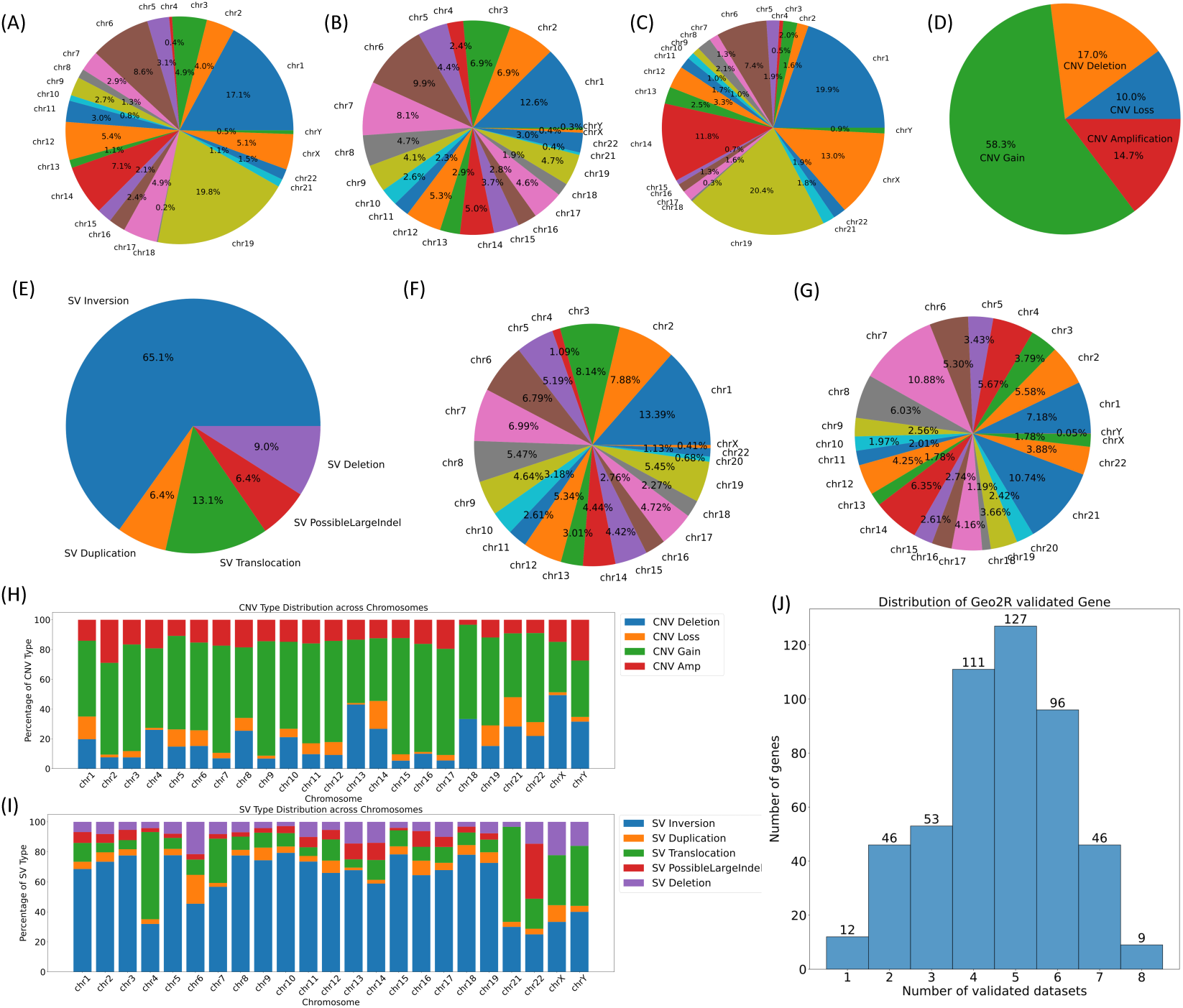
Genomic Aberrations Overview (CNVs, SVs, and LOF) in MM Samples from AIIMS and MMRF Repositories. The figure in panels (A)-(C) displays the chromosome-wise distribution of CNVs, SVs, and LOF. Panel (D) presents the distribution of CNV types identified in MM samples from both AIIMS and MMRF datasets. Similarly, panel (E) shows the distribution of SV types identified in MM samples from the MMRF dataset. Notably, SV analysis was conducted exclusively for MMRF samples due to the absence of WGS data in the AIIMS repository. Continuing SV analysis, panels (F) and (G) exhibit the chromosome-wise distribution of inversions and translocations found in MM samples. Panels (H) and (I) provide the distribution of CNV and SV types for each chromosome individually. (J) Distribution of the top 500 genes validated through MM-related studies using the Geo2R tool. The x-axis represents the number of MM-related studies validating the gene, while the y-axis indicates the count of genes.

### 3.6. Design of 295-gene targeted sequencing panel

To design an effective targeted sequencing panel, we refined the initially identified top-ranked genes based on their significant alterations and the collective impact of their variant profiles in MM. Firstly, we considered four critical variant profiles to identify the candidate driver gene panel: 1. SNV profile, 2. CNV profile, 3. SV profile, and 4. LOF profile. We also integrated the Geo2R validation profile to include MM-relevant genes in the targeted sequencing panel. Finally, we excluded genes that were neither disease-transformative nor disease-initiating. For the SNV profiling of the top 500 significantly altered genes, we filtered based on the median SNV count and the number of samples with non-synonymous SNVs, resulting in 79 genes. The features extracted for SNV profile analysis are detailed in Table-S4, Supplementary File-3. The variant profiling for CNV, SV, and LOF involved filtering genes based on the number of samples exhibiting that particular variant type, yielding 126, 191, and 147 genes, respectively. The features extracted for CNV, SV, and LOF profile analysis can be found in Table-S5, Table-S6, and Table-S7, Supplementary File-3.

By combining genes from SNV, CNV, SV, and LOF variant profiles, we arrived at a comprehensive set of 354 genes. To ensure relevance, we retained genes validated in at least one MM-related study using Geo2R validation. Of the 354 genes, 346 were validated through Geo2R validation analysis. In the final selection, we focused on 212 disease-transformative and 70 disease-initiating genes, resulting in a list of 282 genes. Further, we also included 13 well-established MM biomarker genes that included nine disease-initiating genes (CCND1, CKS1B, IGH, IGK, IGL, MAF, MAFB, MYC, and PRDM1) and four disease-transformative genes (ATM, CD27, HIST1H1E, and JAK1), leading to the design of our proposed 295-gene panel (Table-3, Table-S16, Supplementary File-8). The workflow for designing the 295-gene panel is illustrated in Figure-2.

**Table 3:**
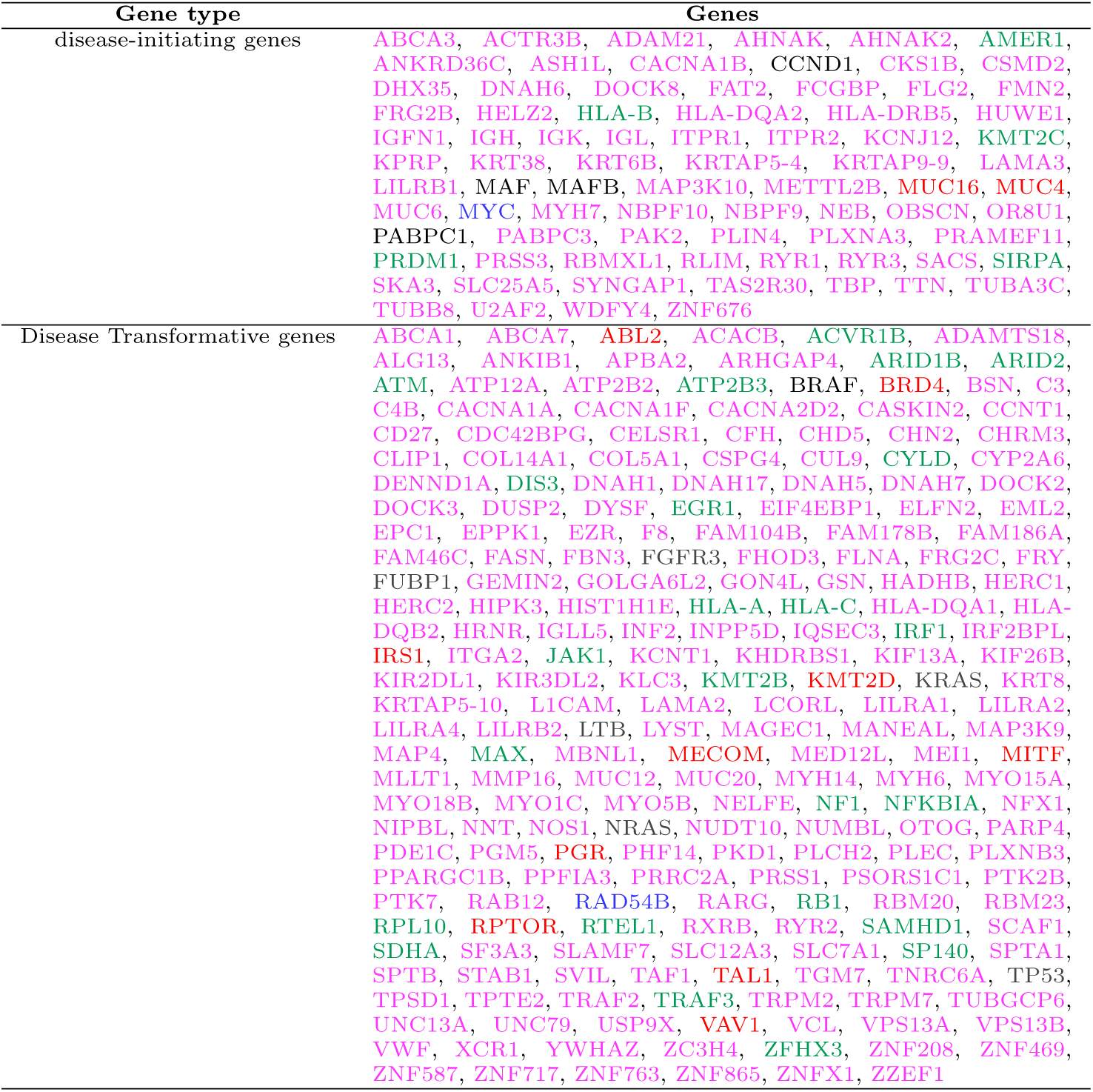
List of disease-transformative and disease-initiating genes in the proposed 295 gene panel. ‘Red’ color denotes oncogenes (OG), ‘black’ denotes genes which are both oncogenes and driver genes (ODG), ‘green’ denotes tumour suppressor genes (TSG), ‘blue’ denotes actionable genes (AG), and ‘magenta’ color is for the rest of the genes which are not previously reported as OG/TSG/ODG/AG in multiple myeloma.

In this panel, four genes, namely, HLA-A, HLA-B, HLA-DRB5, and RYR3, were heavily mutated in all four variant profiles. Additionally, 122 and 32 genes were substantially mutated in at least two and three variant profiles, respectively. Within the MM cohort, notable previously reported significantly altered genes were present including *BRAF, IGLL5, IRF, IGH, MYC, JAK, MAF, KRAS, TP53, TRAF2/3*, among others. Similarly, the MGUS cohort exhibited previously reported genes like *HLA-B, LILRB1, PABPC1, PRSS3*, among others. Several previously reported genes were found in both MM and MGUS cohorts, including *HLA-B, PRSS3, KMT2C*, among others, illustrating their potential role as shared genomic features in the progression from MGUS to MM. We determined each gene’s most prevalent molecular aberration, such as CNV gain, CNV loss, SV translocation, LOF, etc. We observed that CNV gain was the most frequent molecular aberration found in 188 out of the 295 genes, while LOF was the least common, identified in 12 out of the 295 genes. To refine the targeted sequencing regions further, we assessed the most affected coding regions using the UCSC Genome database. The targeted sequencing panel of 295 genes covered 9,417 coding regions in the human genome, spanning a genomic region with a total length of 2.630 Mb in the human genome (Table-S17, Supplementary File-8).

### 3.7. Identification of Significantly altered pathways and ranking of pathway of 295-gene panel

Of the 295-gene panel, only 70 genes were found to be significantly altered in the MGUS cohort, while all 295 genes were part of the MM cohort. We utilized the Enrichr database to identify significantly altered KEGG and Reactome signalling pathways associated with the 295 genes of the MM cohort and those associated with the 70 genes of the MGUS cohort. A total of 39 KEGG and 25 Reactome pathways exhibited significant alterations for the MGUS cohort (see Table-S8 in Supplementary File-4), while 123 KEGG and 50 Reactome pathways were observed to be significantly affected for the MM cohort (refer to Table-S9 in Supplementary File-4). We categorized the significantly altered pathways into four distinct groups according to their significance level changes during the MGUS to MM transition:

Category-1: Pathways increasing in significance in MGUS to MM progression.

Category-2: Pathways decreasing in significance in MGUS to MM transition.

Category-3: Pathways significantly altered in MM but not in MGUS.

Category-4: Pathways significantly altered in MGUS but not in MM.

The complete list of significantly altered pathways for these categories is provided in Tables-S10 and Table-S11 of Supplementary File-4. A total of 32 KEGG and 13 Reactome pathways became more significant as the disease progressed from MGUS to MM, while 5 KEGG pathways and 7 Reactome pathways displayed reduced significance with disease progression from MGUS to MM. A total of 86 KEGG and 30 Reactome pathways were significantly altered only in MM and not in MGUS. Notably, 37 out of 86 KEGG pathways and 5 out of 30 Reactome pathways showed no overlapping genes, with 70 significantly altered genes in MGUS. Lastly, 2 KEGG pathways and 5 Reactome pathways were observed as significantly altered only in MGUS and not in MM.

To determine the top-ranked pathways, we ranked significantly altered pathways in MM based on their adjusted *p*-values (refer to Table-S12, Supplementary File-4). This analysis revealed a selection of MM-related signalling pathways, notably encompassing the antigen processing and presentation, PI3K-AKT signalling pathways, and B-cell receptor prominently featured among the top-ranking pathways.

### 3.8. Analysis of identified gene communities with reference to the 295-gene panel

We employed a five-fold cross-validation training strategy to obtain five distinct learned adjacency matrices for five classifiers, each with a dimension of 798x798. We applied the Leiden algorithm to the respective learned adjacency matrix for each classifier to identify gene communities. Consequently, we derived 5, 5, 6, 5, and 6 gene communities using the learned adjacency matrices from the first, second, third, fourth, and fifth classifiers, respectively. We ranked the communities within each classifier based on the number of previously reported genes present and selected the top three gene communities for each. Subsequently, we merged these top three gene communities for each classifier, resulting in five new distinct learned adjacency matrices with dimensions of 500x500, 500x500, 539x539, 500x500, and 422x422. In the following step, we merged these five distinct newly learned adjacency matrices by computing the mean of gene-gene interactions across the five classifiers. In cases where a specific gene-gene interaction was absent in any fold, we assigned a zero weight for the corresponding interaction in that fold. This process yielded a final adjacency matrix with dimensions of 690 x 690.

Finally, we identified five gene communities from the final learned adjacency matrix using the Leiden algorithm, yielding communities having 202, 125, 122, 104, and 21 genes. The pseudo codes for community detection are provided in Supplementary File-5. The first gene community, comprising of 202 genes, contained 11 OGs, 21 TSGs, 3 ODGs, and 8 AGs. Similarly, the second gene community, with 125 genes, contained 14 OGs, 8 TSGs, 6 ODGs, and 10 AGs. The third gene community, comprising 122 genes, did not include any OGs, TSGs, ODGs, or AGs. The fourth gene community, with 104 genes, contained 4 OGs, 11 TSGs, 1 ODG, and 1 AG. Lastly, the fifth gene community, comprising 21 genes, contained 2 OGs and no TSGs, ODGs, or AGs. The list of genes present in all five gene communities and previously reported genes within each are provided in Table-S13 and Table-S14 of Supplementary File-6. Visualization of all five gene communities, including the top 250 genes and previously reported genes (regardless of their rank), is presented in Figure-6(A)-(E).

**Figure 6:**
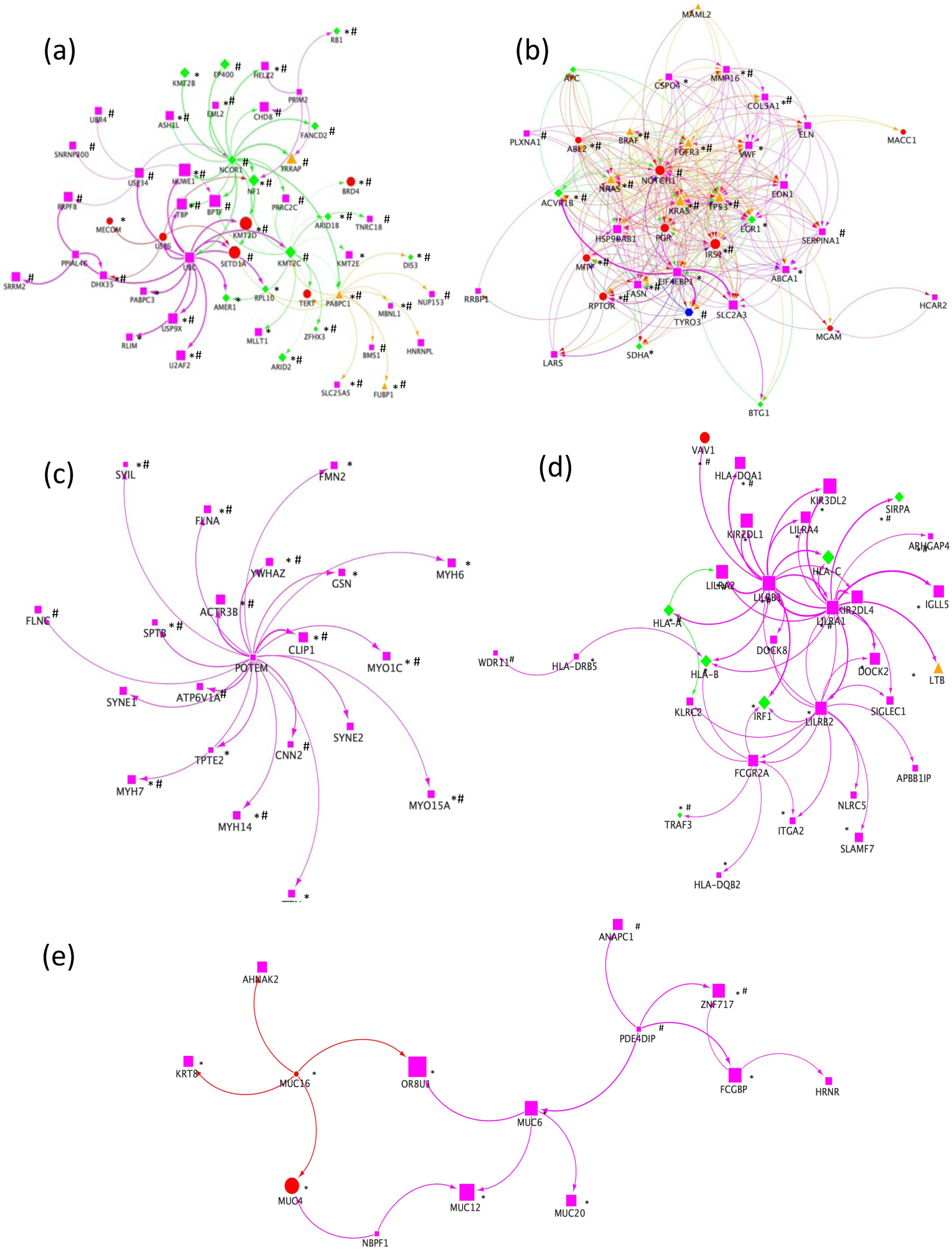
Directed gene community visualization using the learned adjacency matrix obtained from five trained BIO-DGI (PPI9) classifiers. In this figure, (a), (b), (c), (d), and (e) represent the top genes in the first, second, third, fourth and fifth gene communities, respectively. These figures showcase the previously reported genes (OG, TSG, ODG, AG) regardless of their rank, alongside other non-reported genes (in magenta colour) within the top 250 ranks, respectively. Genes marked with “*” are included in the 295-genes panel. Similarly, genes marked with “#” are also highly likely haploinsufficient, with a GHIS score > 0.52. red circle shape node represents oncogene, green diamond shape node represents TSG, orange triangle shape node represents genes that are both oncogene and driver gene, blue hexagon shape node actionable gene and magenta square shape node represents genes that are not previously reported OG/TSG/ODG/AG.

### 3.9. Analysis of CNVs, SVs and LOF associated with 295-gene panel

In addition to analyzing SNV profiles, we comprehensively investigated CNVs, SVs, and LOF in the MM cohort. CNV identification was performed using CNVkit on AIIMS MM samples and on exome segment data from MMRF CoMMpass for MMRF samples. Processed SV data from MMRF CoMMpass was utilized to identify key SVs in MM and 295-genes panel designing. For identifying genes with LOF within a sample, we employed established criteria to evaluate disruptions in gene transcripts due to deletion of essential coding segments, exons, splice signals, or frameshift-inducing deletions [63]. We studied both CNVs and SNVs to identify genes with LOF within each sample. CNVs, SVs, and LOF analysis in the 295 genes revealed crucial molecular aberrations in MM. Chromosome-wise distribution analysis indicated that chr19 (18.03%), chrX (14.2%), and chr1 (9.02%) were notably affected by CNVs (7(A)). Similarly, chr1 (11.4%), chr6 (9.3%), and chr (6.8%) showed prominent SV involvement (7(B)), while chrX (20.43%), chr16 (13.27%), and chr1 (12.52%) were most affected by LOF (7(C)). The majority of CNVs were gains (50.8%) and deletions (16.9%) (7(D)), while inversions (59.2%) and translocations (18.3%) dominated the SV landscape (7(E)). Notable chromosomes impacted by inversion SV included chr1, chr3, chr7, and chr8 (7(F)), and translocations mainly affected chr7, chr11, chr8, and chr1 (7(G)). The distribution of CNV and SV types within each chromosome highlighted their relative abundance (7(H) and 7(I)).

### 3.10. Geo2R validation of the proposed 295-gene panel

We ascertained the relevance of the proposed 295-gene panel with reference to MM via results of the existing MM-related studies through Geo2R validation. Geo2R tool is one of the most widely used tools for identifying significantly dysregulated genes using gene expression or microarray data from previously published studies. We considered 11 MM-related studies for this validation, identifying significantly expressed genes with an adjusted *p*-value of *≤* 0.05 and compared them with our top-ranked genes. Remarkably, out of 295 genes, 268 genes were validated in at least two MM-related studies (Table-S17, Supplementary File-7). Moreover, 68 (23.05%), 77 (26.10%), and 54 (18.30%) genes were found to be significantly dysregulated in MM across datasets related to three, four and five MM-related studies, respectively, as depicted in Figure-7(J).

**Figure 7:**
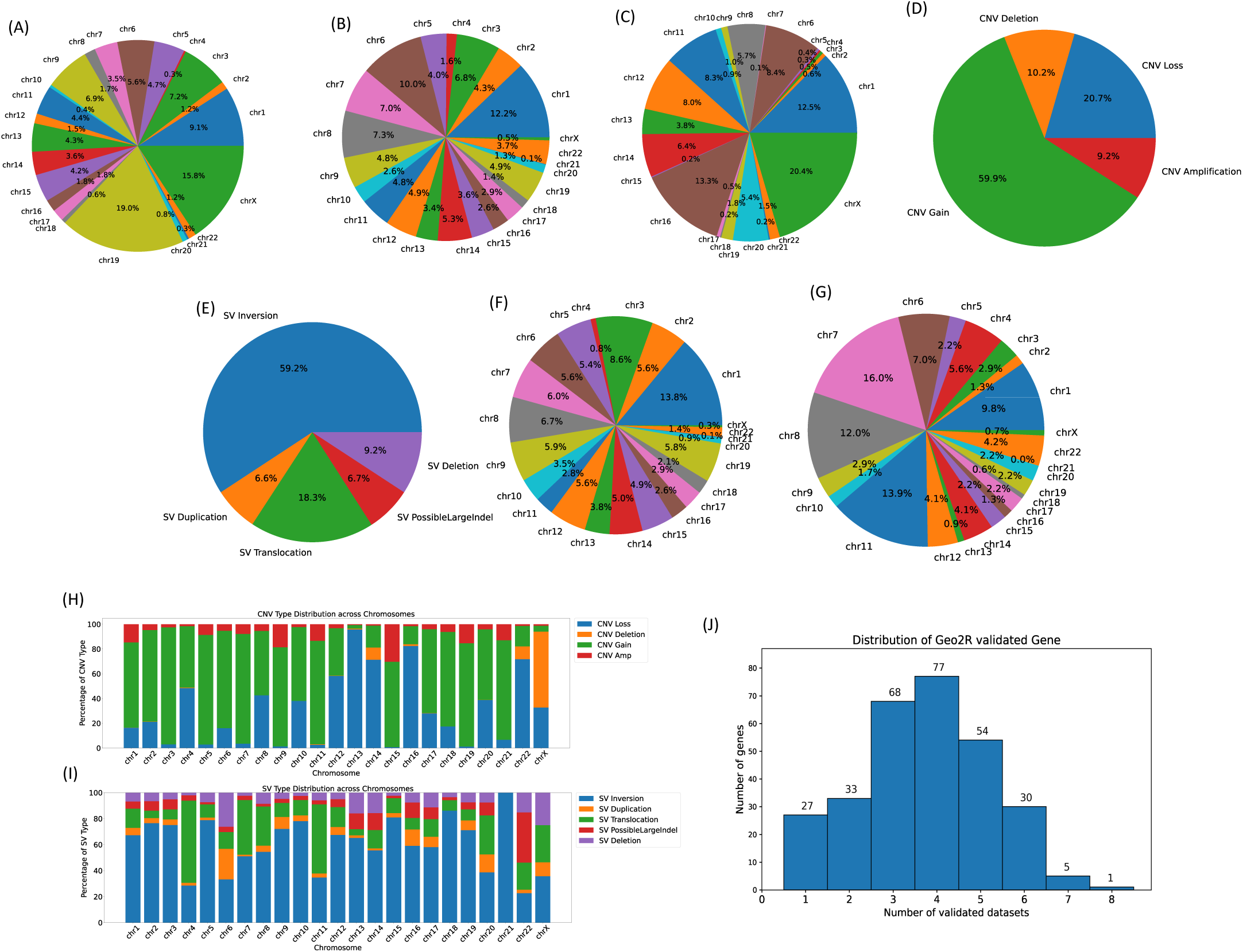
Overview of genomic Aberrations associated with 295 genes (CNVs, SVs, and LOF) in MM Samples from AIIMS and MMRF Repositories. The chromosome-wise distribution of genomic features is shown in panels (A) for CNVs, (B) for SVs, and (C) for LOF. Panel (D) presents the distribution of CNV types identified in MM samples from both AIIMS and MMRF datasets. Similarly, panel (E) shows the distribution of SV types identified in MM samples from the MMRF dataset. Notably, SV analysis was conducted exclusively for MMRF samples due to the absence of WGS data in the AIIMS repository. Continuing SV analysis, panels (F) and (G) exhibit the chromosome-wise distribution of inversions and translocations found in MM samples. Panels (H) and (I) provide the distribution of CNV and SV types for each chromosome individually.(J) Distribution of 295-gene panel validated through MM-related studies using the Geo2R tool. Here, the x-axis represents the number of MM-related studies validating the gene, while the y-axis indicates the count of genes.

### 3.11. Clinical relevance of targeted sequencing 295-gene panel

We performed a two-fold univariate survival analysis on a targeted sequencing panel comprising 295 genes to comprehend how gene variant profiles affect clinical outcomes in MM patients (Figure-8). We utilised two distinct approaches to gauge the effect of gene variant profiles on MM sample clinical outcomes. In the first approach, we individually assessed the impact of each variant profile (SNV, CNV, SV, and LOF) on clinical outcomes using univariate survival analysis. Notably, 168 of the 295 genes significantly influenced clinical outcomes based on at least one variant profile. Of these, 30, 88, 27, and 79 genes significantly impacted clinical outcomes based on SNV, CNV, SV, and LOF variant profiles as prognostic factors, respectively (Table-S16, Supplementary File-8). In the second approach, we amalgamated all four variant profiles into a single feature vector using the FAMD method, leveraging the FAMD first component as a prognostic factor for univariate survival analysis. Subsequently, we found that 188 of the 295 genes significantly influenced clinical outcomes based on the FAMD first component. Combining the clinically relevant genes from the two approaches mentioned above, we discovered that 226 of the 295 genes were clinically relevant for MM. We analyzed the remaining 69 genes that did not show significance in any of the mentioned approaches; we examined them and retained them in the proposed gene panel as these genes were heavily mutated in at least one variant profile (Table-S16, Supplementary File-8). The forest plots and survival curves of genes found significant in at least three variant profiles (e.g. WDFY4, EGR1, IGNF1, INF2, PRSS1, etc.) are shown in Supplementary File-10.

**Figure 8:**
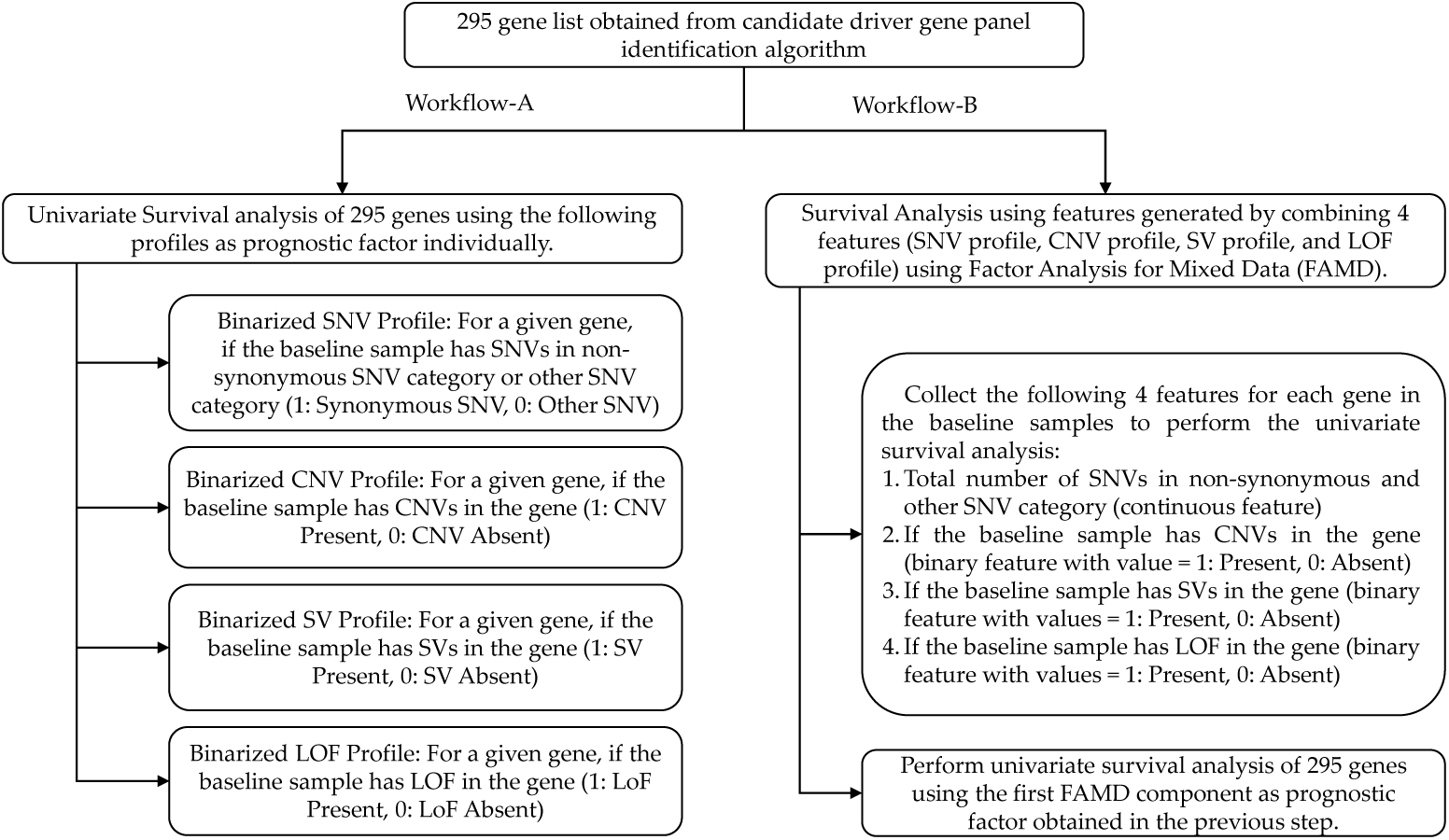
Workflow for two-fold survival analysis of proposed 295-gene panel. In this workflow, we estimated the clinical relevance of gene variant profiles on MM patient clinical outcomes using two distinct approaches. In the first approach (Workflow-A), We individually assessed the impact of each variant profile (SNV, CNV, SV, and LOF) on clinical outcomes. Univariate survival analysis was performed for each variant profile, providing insights into their impact. Using this approach, 167 out of the 295 genes significantly influenced clinical outcomes in univariate survival analysis based on at least one prognostic factor. In the second approach (Workflow-B), we amalgamated the four variant profiles (SNV, CNV, SV, and LOF) for each gene using Factor Analysis for Mixed Data (FAMD), enabling us to estimate a joint feature. Subsequently, we performed univariate survival analysis using the FAMD 1st component as a prognostic factor. In this approach, 187 out of 295 genes demonstrated significance in univariate survival analysis based on the FAMD 1st component (a combined feature generated by integrating the four variant profiles for each gene). Interestingly, 129 genes out of these 173 were also identified as significant in univariate survival analysis using Workflow-A. By combining both approaches, out of 295, 225 genes were found to influence the clinical outcomes of MM patients significantly.

## 4. Discussion

Multiple Myeloma (MM) is a malignancy that typically progresses from premalignant stages, often starting with MGUS [84]. A targeted sequencing panel is important for the precise characterization of the genomic alterations to understand the risk of progression, enabling timely interventions and ultimately improving patient outcomes. Recent studies have shed light on the genomic events that drive the transformation from premalignant stages to MM [85, 86, 87, 88]. Moreover, a number of studies have proposed targeted sequencing panels for molecular profiling of MM patients based on previously identified genomic events in MM and MGUS [17, 18, 19, 20, 21]. However, none of these studies have taken into account the design of the panel using biomarkers and gene-gene interactions that have the potential to distinguish MM from MGUS.

In this study, we addressed this challenge by designing a targeted sequencing panel of 295 genes hosting key genomic biomarkers. We designed an AI-powered attention-based bio-inspired BIO-DGI (PPI9) model to identify the key genomic biomarkers and gene interactions for panel crafting. The BIO-DGI (PPI9) model is biologically inspired, learning to identify distinguishing patterns of MM and MGUS using gene-gene interactions and their corresponding genomic features. Genes with a higher number of interactions are deemed more biologically relevant. We specifically considered deleterious SNVs associated with MM and MGUS, resulting in highly MM-relevant, significantly altered genes being ranked at the top. The inclusivity of three global repositories having MM and MGUS cohorts with diverse ethnicities, the ability of the AI-based workflow to comprehend gene inter-dependencies, extensive benchmarking, and rigorous post-hoc analysis collectively render the BIO-DGI (PPI9) model innovative and highly efficient.

During classification, the functional significance of nonsynonymous SNVs, as quantified by Phylop scores, emerged as the most prominent genomic feature. Following closely, the allele depth of synonymous SNVs and the overall count of other SNVs (encompassing non-frameshift insertions/deletions/substitutions, intronic, intergenic, ncRNA intronic, upstream, downstream, unknown, and ncRNA splicing SNVs) ranked as the second and third most influential genomic features, respectively (Figure-4). These findings are in line with the literature because the impact of synonymous SNVs across various cancer types has been highlighted by various studies [89, 90, 91, 92, 93]. In the post-hoc analysis for model interpretability, we utilized the ShAP algorithm to identify the top-ranked genes within the top-performing models. Table-1A, Table-1B, Table-2 provides an overview of the total number of previously reported genes present in the 798 significantly altered genes and those identified by top-performing models, presenting complete gene lists under top-250 and top 500 ranks. Notably, the BIO-DGI (PPI9) model outperformed by identifying the highest number of previously reported genes, encompassing known oncogenes (OGs) such as BIRC6, MUC4, NOTCH1, PGR, SETD1A, and VAV1, tumour suppressor genes (TSGs) like DIS3, EP400, MYH11, SDHA, both oncogenes and driver genes (ODGs) such as KRAS, NRAS, TP53, TRRAP, and actionable genes (AGs) including APC, ARID1B, MITF, NFKBIA, and TYRO3. Interestingly, most of these genes (except ODGs) display high relevance to MM despite not being explicitly reported as MM driver genes.

Additionally, our analysis identified MUC6, LILRA1, and LILRB1 as the top three genes contributing significantly to the classification of MM and MGUS, although none of these have been previously categorized as OGs, TSGs, ODGs, or AGs in the literature. Here, this is to note that MUC6 gene is associated with the immune system pathway, playing a crucial role in MM development and progression [94]. Similarly, the other two genes, LILRA1 and LILRB1, are associated with the innate immune system pathway. Notably, LILRB1 has been reported to be associated with MM pathogenesis as an inhibitory immune checkpoint for B-cell function in prior studies [95, 96]. We also employed the Geo2R tool to validate the top-ranked genes obtained from the post-hoc analysis of the BIO-DGI (PPI9) model. We included 11 MM-related studies for validation and observed that 488 out of 500 genes were found to be disrupted in MM. This finding ensures the relevance of top-ranked genes in MM.

We curated a 295-gene panel by rigorously analysing variant profiles (SNVs, CNVs, SVs, and LOF) of the top 500 genes. We specifically considered MM-relevant genes that were disrupted in at least one previously published MM study. To identify pivotal genomic events responsible for MM development and progression, we categorized them into different groups based on their occurrence at specific disease stages (MM or both MM and MGUS). Genomic events observed in both MGUS and MM such as translocations associated with the IGH and MYC genes [86, 13, 97, 98, 99, 100] and amp(1q) [13] were labeled by us as “disease-initiating” events, while those that were observed to be present in MM but not in MGUS including del(13q), del(16q), del(17p), etc. [13] were labeled by us as “disease-transformative” events, and are shown in Table-4A and Table-4B.

**Table 4:**
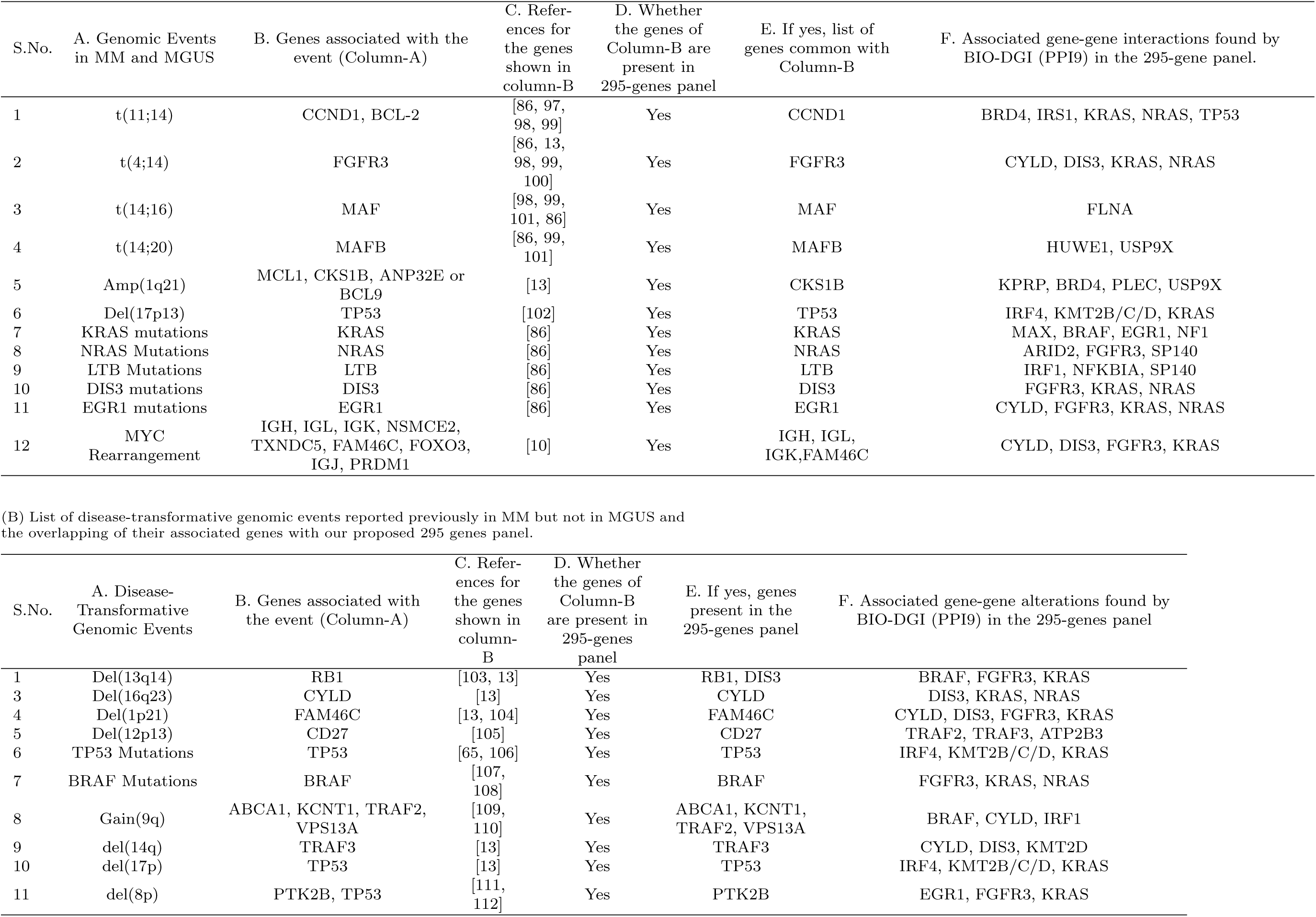
List of key genomic events in MM and MGUS and overlapping of their associated genes with 295 genes panel.

We comprehensively analysed CNVs, SVs, and LOFs identified in the 295 genes panel across multiple myeloma (MM) samples obtained from both AIIMS and MMRF datasets. Our analysis highlighted chr1, chr14, chr19, and chrX as the most affected chromosomes, displaying various CNV genomic alterations. Notably, chr1 exhibited significant alterations, such as amp(1q), associated with disease aggressiveness [13, 113], and del(1p), frequently observed in MGUS [13, 104]. Furthermore, chr14 revealed prevalent translocations involving IGH, such as t(4;14), t(14;16), t(14;20), established as biomarkers in MM [13]. Additionally, CNVs linked to chr19, such as gain(19p) and gain(19q), were significantly more prevalent in MM than in MGUS [84]. Recently, it was shown that abnormalities of chromosome X and MAGE-C1/CT7 expression are much more frequent events in MM than previously reported [114]. The intricate interplay between alterations in these chromosomes and other genetic events contributes to increased genomic instability, facilitating the acquisition of additional mutations that promote MM aggressiveness [11].

Upon scrutinizing the clinical significance of the proposed panel consisting of 295 genes through survival analysis across five variant profiles (SNV, CNV, SV, LOF, and FAMD 1st component of amalgamation of four variant profiles), seven genes (*ACACB, ARHGAP4, ASKIN2, FAM186A, IGFN1, NBPF9*, and *TPTE2*) demonstrated clinical significance in at least four variant profiles. Notably, *ACACB* and *TPTE2* were identified as vulnerable genes in Multiple Myeloma Cells through RNA Interference Lethality Screening of the Druggable Genome [115]. *ACACB* might have been playing a pivotal role in Multiple Myeloma (MM) progression because its top transcription factor binding sites such as AP-1, C/EBPalpha, MAZR, RFX1, STAT1, STAT1*α*, and STAT1*β* play a role in either cell proliferation, differentiation, apoptosis, oncogenesis or in regulating the immune response. For example, the role of MAZR in cell proliferation, apoptosis, and tumorigenesis may indicate its contribution to MM progression by affecting ACACB regulation. Additionally, RFX1, influencing the cell cycle and immune response, could also play a role in ACACB modulation within the context of MM. Lastly, the engagement of STAT1, along with its isoforms STAT1*α* and STAT1*β* in immune response and tumour suppression might implicate ACACB in MM pathogenesis, potentially linking aberrant fatty acid metabolism to the dysregulated immune responses characteristic of the disease. The intricate interplay between ACACB and these transcription factors underscores its multifaceted involvement in MM progression.

Similarly, *TPTE2* ’s involvement in diverse cellular processes, including immune responses, inflammation, and cell survival, critical aspects of MM progression, is suggested by its association with NF-kappaB and its subunits (*NF-kappaB1* and *NF-kappaB2*). Moreover, RelA binding sites signify *TPTE2* ’s involvement in the activation of gene expression in response to stimuli. Altogether, *TPTE2* ’s engagement with these transcription factors indicates its intricate participation in diverse cellular processes, potentially contributing to the complex pathogenesis of multiple myeloma. Further investigations are warranted to elucidate the precise molecular mechanisms and implications of *TPTE2* in MM progression.

We thoroughly evaluated our proposed 295-gene panel, comparing it with five previously published targeted sequencing panels used for MM genomic profiling. These panels were thoughtfully crafted based on MM-related literature and underwent validation using diverse methods such as FISH and analysis of whole-genome sequencing (WGS) data, etc. Upon analyzing the validated variant profiles, we noted that, alongside our proposed panel, Sudha et al. [18] also validated their panel on SNVs, CNVs, and SVs, encompassing translocations linked to IGH and MYC. However, Sudha et al.’s panel validation was carried out on WGS cohorts of MM samples and MM cell lines and did not account for potentially distinguishing genomic biomarkers of MGUS and MM. Furthermore, none of the targeted panels developed so far analyzed the clinical significance of individual genes in their panels on survival outcomes. The present study is unique and adds valuable information on the potential impact of these genes on clinical outcomes in MM. Thus, the proposed panel is unique in that it not only helps identify transforming events in patients with MGUS, but also is powered to assess genomic features that impact treatment outcomes.

Moreover, our panel incorporated MM-relevant genes exhibiting loss-of-function (LOF), a critical consideration lacking in previous panels. Comparing the genes across the previously published panels, we found that 19 out of 47 (34%) genes were common with Kortum et al.’s, 26 out of 182 (10.43%) with Bolli et al.’s, 47 out of 465 (8.38%) with White et al.’s, 18 out of 26 (57.69%) with Cutler et al.’s, and 42 out of 228 (14.5%) with Sudha et al.’s panels, respectively. The comprehensive gene list encompassing all genes from the five panels is provided in Table-S18, Supplementary File-9. Additionally, a detailed comparison of these panels is presented in Table-5 and Table-S19, Supplementary File-9.

**Table 5:**
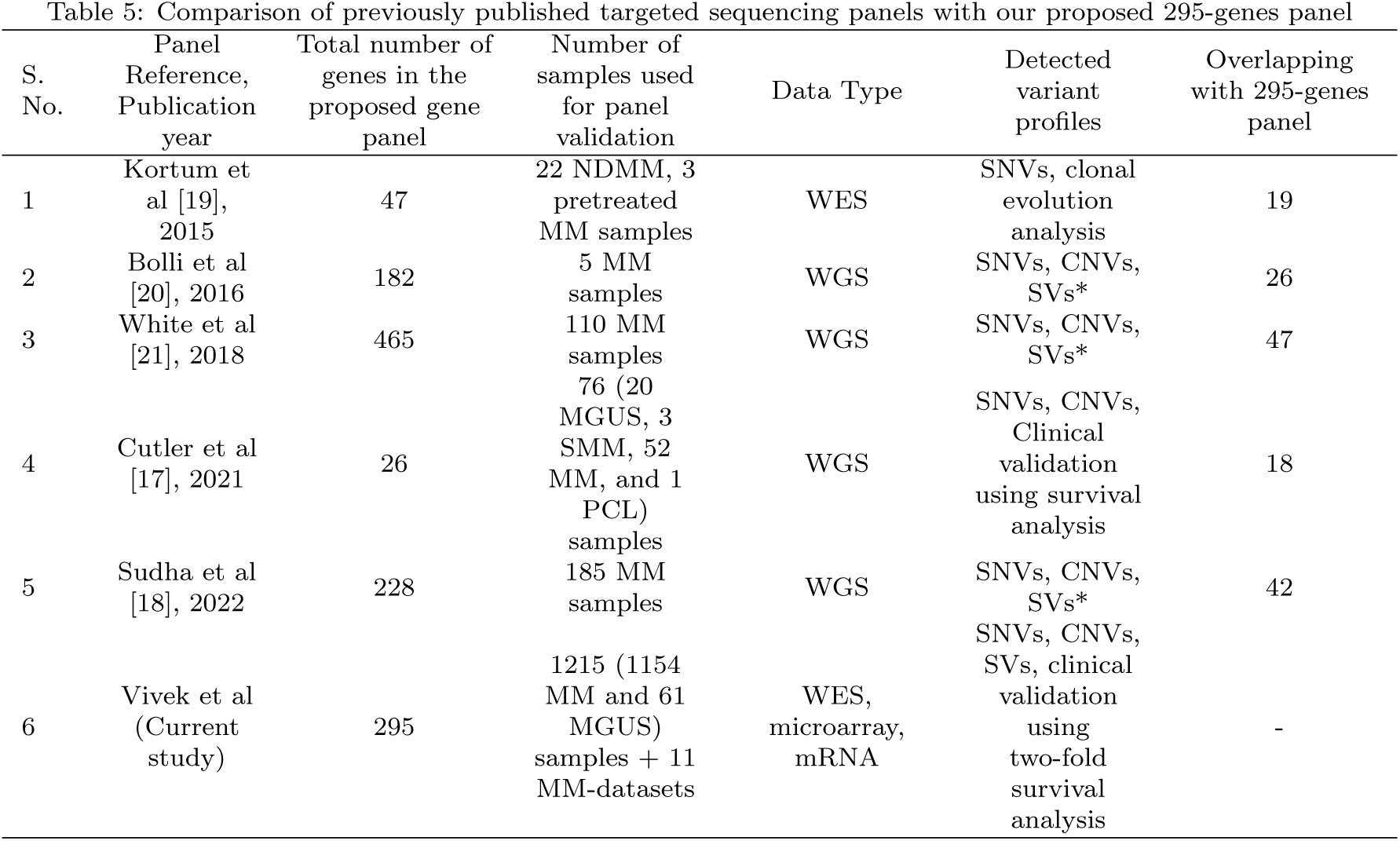
Comparison of previously published targeted sequencing panels with our proposed 295-genes panel.

In addition, we conducted pathway analysis using the Enrichr database to elucidate the significantly altered pathways associated with the 295-gene panel. These pathways were subsequently ranked based on their statistical significance (adjusted *p*-value) to identify the top pathways exhibiting substantial alterations. Notably, a distinct pattern emerged when assessing the significance of altered pathways in relation to disease progression. Pathways associated with various cellular processes displayed significant alterations in MGUS, but their significance diminished as the disease transitioned from MGUS to MM. In contrast, pathways specifically linked to multiple myeloma exhibited pronounced alterations as the disease advanced (Table-S10, S11, Supplementary File-4). Out of the 295 genes, 174 were implicated in significantly altered pathways. Key MM-related pathways, including MAPK signaling, PI3K-AKT, B-cell receptor, Human papillomavirus, and immune system pathways, prominently featured among the significantly altered pathways. The top 50 pathways associated with the proposed 295-gene panel, along with the number of significantly altered genes and their respective rankings for each pathway, are illustrated in the bubble plot presented in Figure-9. These compelling findings warrant further investigation to determine whether the significantly altered genes associated with these pathways could potentially serve as valuable biomarkers of the development of MM during the early stages of the disease, particularly MGUS.

**Figure 9:**
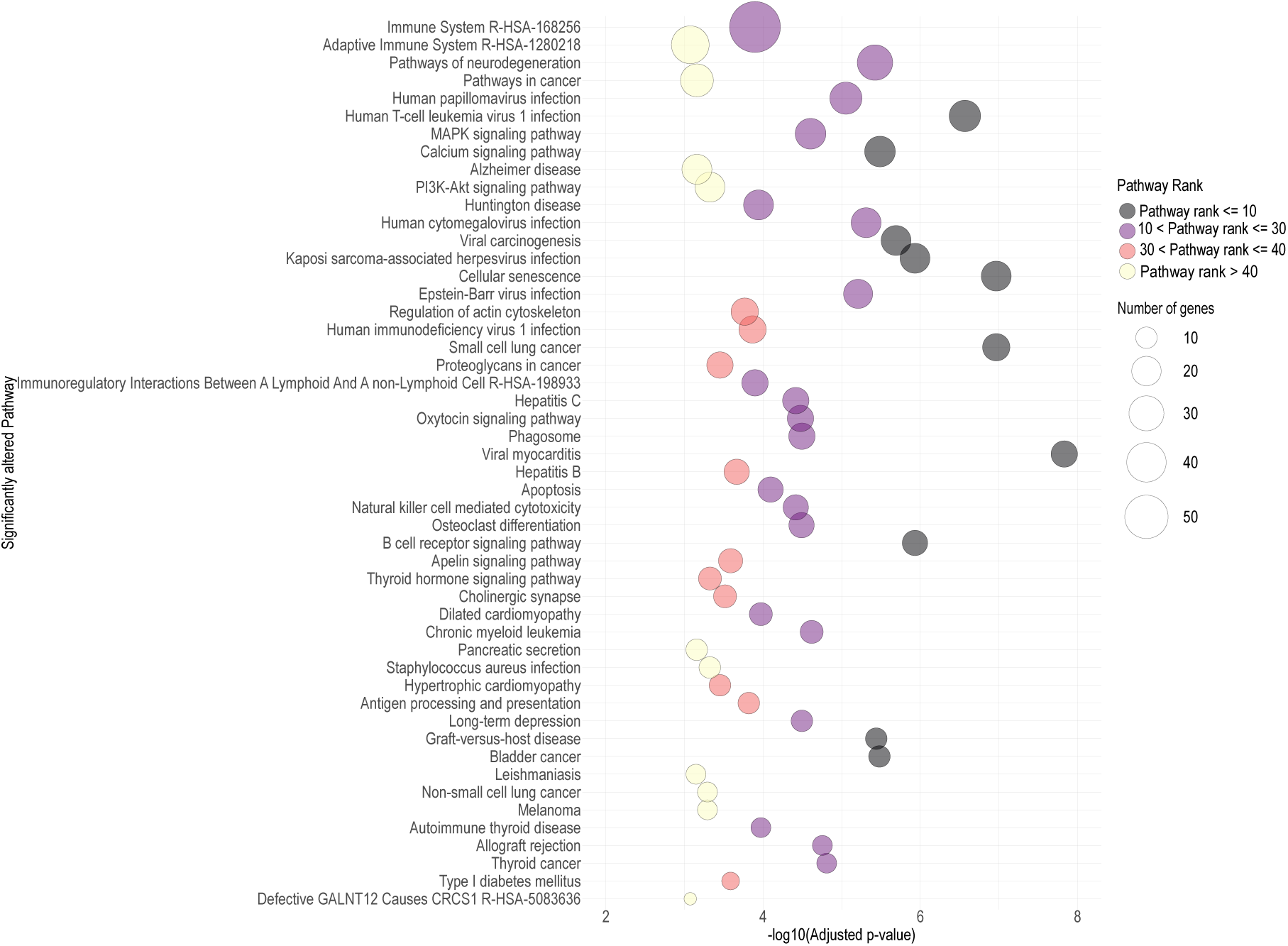
Bubble Plot illustrating the Top 50 significantly altered signalling pathways in MM associated with genes included in the proposed 295-gene panel. The bubble size indicates the number of significantly altered genes linked to the 295-gene panel, and colour signifies the pathway rank. The x-axis represents the -log10 (adjusted *p*-value) score of the pathway, and the y-axis displays the pathway names.

Using the interaction weights acquired from the BIO-DGI (PPI9) model, we identified five gene communities and to enhance the information for each node within a gene community, we integrated node influence determined by the Katz centrality score and likelihood of haploinsufficiency gauged through the GHIS score. The genes surpassing the median GHIS score of 0.52 (Figure 6) notably include, UBC, USP6, PRIM2, USP34, KMT2C, PABPC1, and NCOR1 in the first gene community (Figure-6(A)), TP53, NRAS, IRS1, EIF4EBP1, HSP90AB1, and FGFR3 in the second gene community (Figure-6(B)), POTEM in the third gene community (Figure-6(C)), and LILRA1, LILRB1, LILRB2, FCGR2A in the fourth gene community (Figure 6(D)) and appear as central genes that shows that these might have been playing a significant role in MM pathogenesis. This is to note that these genes are already known to be associated with MM. Furthermore, we observed that, out of the 295 genes, 67 displayed substantial node influence within the gene community, encompassing various previously reported MM-relevant genes like *BRAF, HLA-A/B, FGFR, IRS1, NRAS*, and *SDHA*. Additionally, 74 genes exhibited a high likelihood of haploinsufficiency, including several previously reported MM-relevant genes such as *ARID1B, FGFR, NRAS, TRAF2*, and *ZNF717*. Moreover, 32 genes displayed both high node influence and a high likelihood of haploinsufficiency, including *FGFR, HUWE1, KRAS, KMT2C/D, TP53*, and *ZNF717*. We strongly recommend further analysis on these central genes to unveil their role in disease progression.

While examining the gene communities and their involvement in key genomic events of MM, we noted several genes with substantial node influence and likelihood of actively participating in these events. For instance, in the first gene community (Figure-6(A)), seven genes (*BRD4, DIS3, HUWE1, RB1, SLC25A5, RB1*, and *USP9X*) were associated with genomic events observed in both MM and MGUS. Similarly, the second gene community (Figure-6(B)) included five genes (*EGR1, IRS1, KRAS, NRAS,* and *TP53*) involved in genomic events observed in both MM and MGUS. In the third gene community, *FLNA* was found to be associated with genomic events observed in both MM and MGUS. The fourth community featured *LTB* associated with genomic events observed in both MM and MGUS, while *TRAF3* was associated with genomic events observed in MM only. Finally, the fifth gene community had no genes linked to the key genomic events shown in Table-4A and Table-4B. The presence of genes actively participating in MM-related key genomic events, displaying high node influence within the community, and a high likelihood of haploinsufficiency underscores the relevance of our proposed targeted sequencing panel in MM and MGUS.

In this study, we investigated the interactions of the 295-gene panel with drugs by leveraging the DGIdb database [116]. Our focus was on identifying drugs associated with these genes based on the strength of evidence for interaction, considering factors such as the number of publications and sources supporting each claim. The top 15 drugs with the most robust gene interactions were prioritized, involving well-known genes like *BRAF, KRAS, FGFR, TP53*, among others. The resulting gene-drug interaction network, depicted in Figure 10, was both weighted and directed. Among the top three drugs demonstrating interactions with key driver genes in multiple myeloma were Dabrafenib, Trametinib, and Vemurafenib. Notably, Vemurafenib and Dabrafenib are recognized as BRAF inhibitors (BRAFi) [117] and have been considered in the context of multiple myeloma treatment [118, 119]. Additionally, Cisplatin, a drug known for its inclusion in the potent combination therapy VTD-PACE (bortezomib-thalidomide-dexamethasone-cisplatin-doxorubicin-cyclophosphamide-etoposide) [120], demonstrated significant interactions. Furthermore, our analysis highlighted drugs commonly used in the treatment of other cancers, such as Carboplatin, Docetaxel, and Fluorouracil, showing interactions with key driver genes in multiple myeloma. These findings suggest the potential of these drugs as novel chemotherapeutic agents for multiple myeloma.

**Figure 10:**
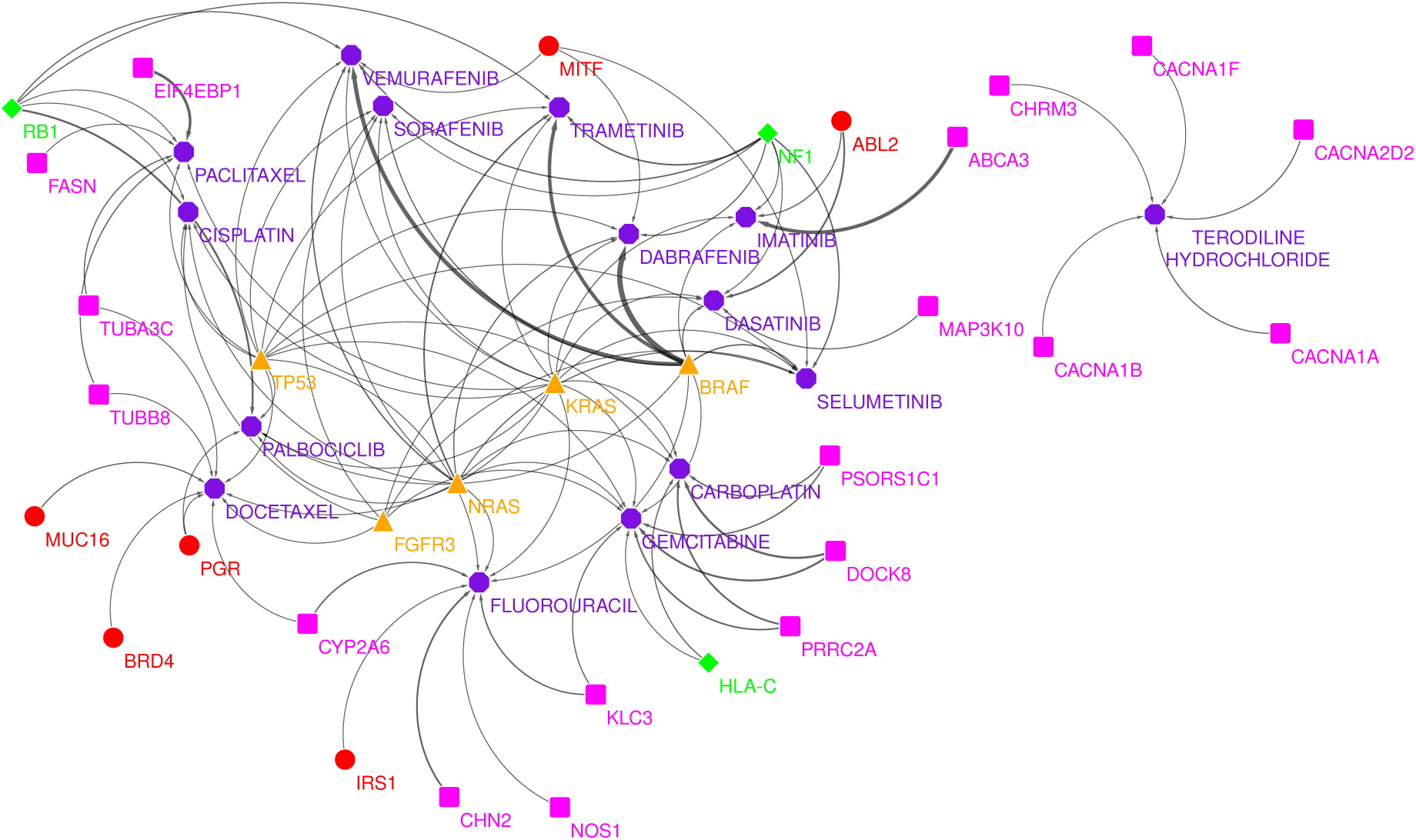
Gene-drug interactions for genes in 295 genes panel. In the above network, the colour and shape notations are as follows: red circle represent oncogenes, orange triangle represent genes that are both oncogenes and driver genes (ODG), magenta square represent genes that are not previously reported as OG/TSG/ODG/AG, purple octagon represent drugs. The edge width represents the mean of gene-drug interaction scores obtained from DGIdb 4.0 database [116].

## 5. Conclusions

Distinguishing Multiple Myeloma from its precursor stage, MGUS, and identifying those at risk of progression to overt MM poses a significant challenge due to overlapping genomic characteristics. The present study addresses this challenge by leveraging the innovative AI-based BIO-DGI (PPI9) model, incorporating gene interactions from nine PPI databases and exonic mutational profiles from global MM and MGUS repositories (AIIMS, EGA, and MMRF). Our study demonstrates superior quantitative and qualitative performance with application-aware interpretability. The model identified a substantial number of previously reported genes, including Oncogenes (OGs), Tumor Suppressor Genes (TSGs), Oncogene Databases (ODGs), and Associated Genes (AGs), known for their high relevance in MM. Geo2R validation of the 295-gene panel, coupled with an association with MM-relevant pathways, underscores the panel’s pertinence to MM. Our analysis highlights the functional significance of non-synonymous mutations, allele depth of synonymous SNVs, and the total number of other SNVs as crucial genomic biomarkers in distinguishing MM from MGUS. Significant alterations in chromosomes 1, 14, 19, and X suggest the inclusion of genes associated with these chromosomes in genomic evaluation of myeloma patients. Within this panel, we identified genes with substantial node influence and prominent gene-gene interactions from five gene communities, shedding light on crucial gene biomarkers and their interactions pivotal to MM pathogenesis. These findings hold immense potential for informed therapeutic interventions, facilitating early detection and interception of disease progression in MM. The proposed panel, driven by innovative AI modelling and comprehensive genomic analysis, emerges as a promising tool for advancing our understanding of MM and improving patient outcomes through precision medicine.

## Acknowledgements

This work was supported by a grant from the Department of Biotechnology, Govt. of India [Grant: BT/MED/30/SP11006/2015] and the Department of Science and Technology, Govt. of India [Grant: DST/ICPS/CPS-Individual/2018/279(G)]. Authors acknowledge dbGaP (Project#18964) for providing authorized access to the MM datasets (phs000748 and phs000348). The Multiple Myeloma Research Foundation provided funding support for the phs000348 study in collaboration with the Multiple Myeloma Research Consortium. The Broad Institute Genome Sequencing, Genetic Analysis, and Biological Samples Platforms provided assistance with data generation, processing and analysis. The datasets used for the analyses described in this work were obtained from dbGaP through dbGaP accession number phs000348.v1.p1. Data of study phs000748 were generated as part of the Multiple Myeloma Research Foundation CoMMpass [SM] (Relating Clinical Outcomes in MM to Personal Assessment of Genetic Profile) study (www.themmrf.org). We also acknowledge EGA (EGAD00001001901) for providing authorized access to the MGUS data. The authors would also like to thank the Centre of Excellence in Healthcare, IIIT-Delhi, for their support in this research. Authors acknowledge the valuable insights provided by Dr Satish Sankaran, Dr. Nandini Pal Basak, and Dr. Mohit Malhotra from Farcast Biosciences Pvt Ltd, India, that greatly improved the gene panel designing workflow.

