## Supplementary File-2 for "A Comprehensive Targeted Panel of 295 Genes: Unveiling Key Disease Initiating and Transformative Biomarkers in Multiple Myeloma"

**Supplementary File-2: (A) AI-based workflow to infer key genomic biomarkers and gene interactions to distinguish MGUS and MM using the whole-exome sequencing (WES) data.**

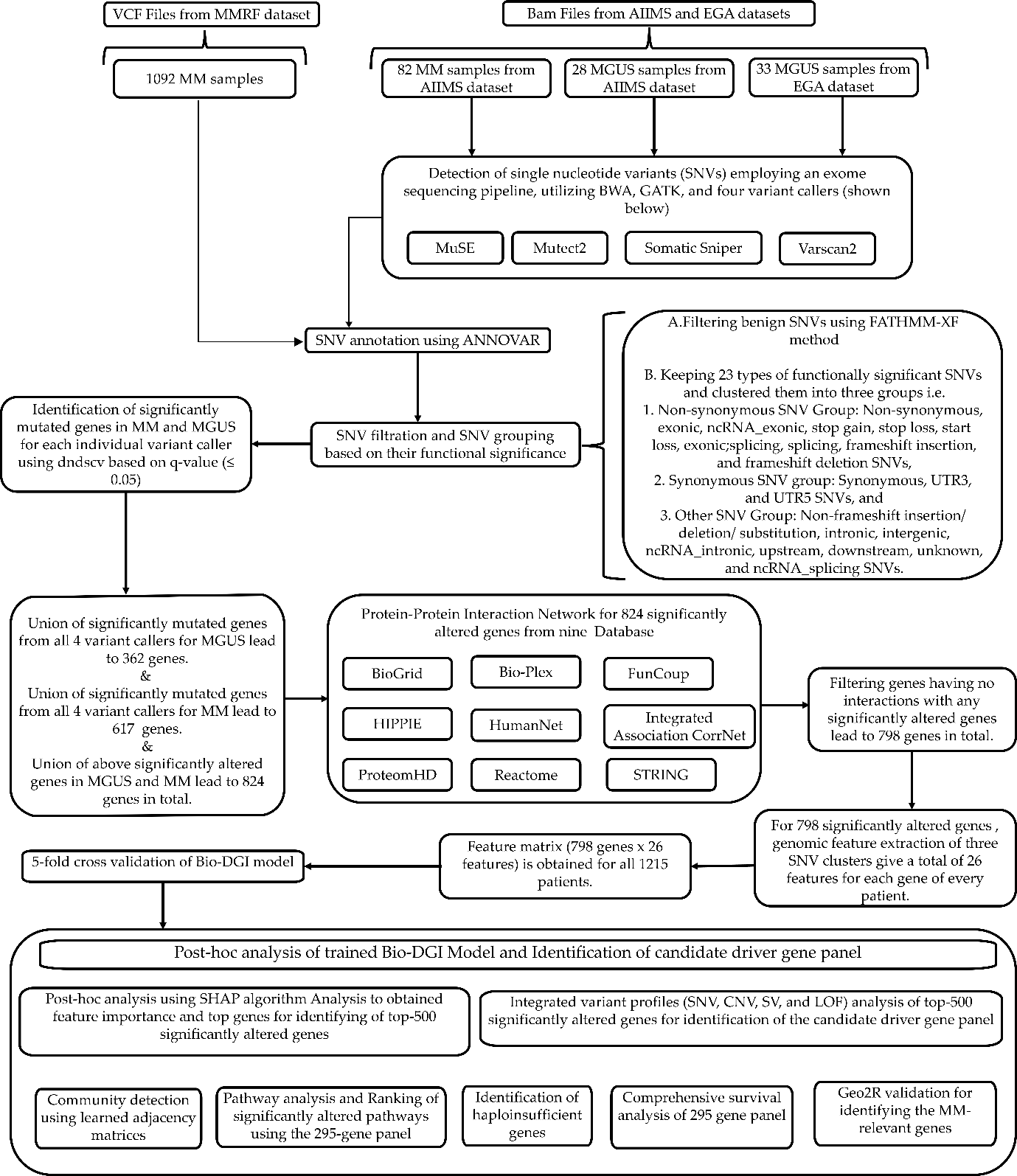

**Supplementary File-2: (B)Architecture and mathematical description of proposed Bio-DGI (PPI9) model**

1. **Architecture of proposed Bio-DGI (PPI9) model:**

Table 1: Hyperparameters values and layer dimensions of the Bio-DGI (PPI9) model architecture

| GCN architecture attribute/ hyperparameter | Hyperparameter value |
| --- | --- |
| No. of GCN layers | 1 |
| GCN layer dimensions | Input sample dimension: 798x26  1st Layer (For each node): 26x1  Output dimension: 798x1 |
| Multi-head attention Units | Number of multi-head attention units= 1 |
| Output linear layer dimension | 798x2 (number of classes = 2) |
| Activation function | LeakyReLU(0.1) |
| Dropout | 0.80 |
| Cost function and adjusted cost for class imbalance | Cost function: NLL Loss  Cost adjusted: 20.0 |
| GCN Weight Initialization | Normal Xavier |

1. **Mathematical description of proposed Bio-DGI (PPI9) model:**

The proposed Bio-DGI (PPI9) model described below integrates Graph Convolution Layers (GCN), Attention Layers, and Attention Consensus Layers for a comprehensive understanding of graph-structured data for identifying the key biomarkers and gene interactions that can differentiate MM from MGUS. The details of the proposed models are as follows:

*A. Data type required for model training:*

The proposed Bio-DGI model is designed to work with graph-structured data, where nodes represent significantly altered genes, and edges represent gene-gene interactions obtained after merging interactions from nine PPI databases.

*B. Layers in proposed Bio-DGI (PPI9) model:*

(i) Graph Convolution Layers (GCN): The Graph Convolutional Networks (GCNs) are utilized to capture the node-level representations by aggregating information from neighbouring nodes in the graph. The GCN layer processes the graph's adjacency matrix and node features to generate initial node embeddings.

Let's say, $A$ as the adjacency matrix of the graph (indicating connections between nodes) of dimension ($798\times798$). $X$ as the feature matrix (containing features associated with each node) of dimension ($798\times26$).

The graph convolution operation in a single layer can be defined as:

$$H^{\left( l+1 \right)}=\sigma\left( \tilde{D}^{\frac{-1}{2}}\tilde{A}\tilde{D}^{\frac{-1}{2}}H^{\left( l \right)}W^{\left( l \right)} \right)$$

where

$H^{\left( l \right)}$ is the output matrix of node representations at layer $l$.

$\tilde{A}=A=I$ is the adjacency matrix with added self-connections.

$\tilde{D}$ is the degree matrix of $\tilde{A}$.

$W^{\left( l \right)}$ is the weight matrix for layer $l$.

$\sigma$ is an activation function.

(ii) Multi-head attention layer: Multi-head attention is a mechanism that enables the model to focus on different parts of the input differently, attending to multiple aspects or features simultaneously. In the multi-head attention mechanism, we have $h$ parallel self-attention operations, each with its own set of learned parameters ($W_{i}$). The outputs of these attention operations are concatenated and linearly transformed to obtain the final output.

Mathematically, the multi-head attention can be expressed as follows:

$$MultiHead\left( Q,K,V \right)=Concat\left( {head}_{1},\ldots..,{head}_{h} \right)W_{O}$$

where each head operates as:

$${head}_{i}=Attention\left( {QW}_{Qi},{KW}_{Ki},{VW}_{Vi} \right)$$

$$Attention\left( Q,K,V \right)=softmax\left( \frac{QK^{T}}{\sqrt{d_{k}}} \right)$$

Where $\left( Q,K,V \right)$ are the input matrices representing queries, keys, and values respectively. $W_{Qi},W_{Ki},W_{Vi}$ are learned weight matrices for each head. $W_{O}$ is the weight matrix for the output projection.

The overall model would integrate these components in a way that leverages the graph structure through GCNs and the attention mechanism to focus on different features via multi-head attention, providing a powerful tool for processing graph-structured data.
