## Supplementary File-5 for "A Comprehensive Targeted Panel of 295 Genes: Unveiling Key Disease Initiating and Transformative Biomarkers in Multiple Myeloma"

Supplementary Material-5: Algorithm for gene community identification using the learned adjacency matrices obtained from five trained Bio-DGI (PPI9) classifiers.

| Step-1 | Fetch learned adjacency matrices for each fold | Fetch the learned adjacency matrix from each of the five Bio-DGI (PPI9) classifiers. | All five learned adjacency matrices are of dimension 798x798. |
| --- | --- | --- | --- |
| Step-2 | Community detection for each fold | - For each of the five learned adjacency matrices, apply Leiden algorithm (LA) for community detection. - Within each fold, rank the communities based on the number of previously reported genes (OG, TSG, ODG, and AG) found in it. | In each of the five-fold learned adjacency matrices, we identified 5 communities in the first fold, 5 communities in the second, 6 communities in the third, 5 communities in the fourth, and 6 communities in the fifth fold learned adjacency matrix. |
| Step-3 | Gene selection from top-communities for each fold | - For each fold, select the top three communities based on the number of previously reported genes (OG, TSG, ODG, and AG) present in the community. - For each fold, generate a new learned adjacency matrix by taking union of the genes present in top three communities. | The union of genes derived from the top three communities in each of the fold-1, fold-2, fold-3, fold-4, and fold-5 learned adjacency matrices resulted in gene sets comprising 500, 500, 539, 500, and 422 genes, respectively. Consequently, using the same gene sets in each of the five folds, we generated new learned adjacency matrices with dimensions of 500x500, 500x500, 539x539, 500x500, and 422x422, respectively. |
| Step-4 | Generation of a consensus of learned adjacency matrix | - Merge all five of the new adjacency matrices obtained in the previous step (achieved by combining genes from the top three communities in each fold) by computing the mean of gene-gene interaction weights across these learned adjacency matrices. - In case, if a particular gene-gene interaction is absent in any fold, assume a weight of zero for the corresponding interaction in that fold. | Upon merging the five new learned adjacency matrices, we obtained a consolidated adjacency matrix with dimensions of 690x690. |
| Step-5 | Community detection on merged adjacency matrix | On the merged adjacency matrix obtained from Step-4, apply LA for community detection using the new gene-gene interaction weights. | A total of five communities were identified with community size 202, 125, 122, 104, and 21. |
